# A conserved archaeal ribosome-associated factor linking bacterial hibernation and eukaryotic energy sensing

**DOI:** 10.64898/2026.02.22.707292

**Authors:** Diorge P. Souza, Mira B. May, Jackson Carrion, Vikram Alva, Alex Bisson, Joseph H. Davis

## Abstract

Ribosome hibernation helps cells survive stress by reversibly silencing translation and limiting degradation of ribosomal complexes. Although well characterized in bacteria and eukaryotes, archaeal hibernation remains poorly understood. Using cryoEM to analyze lysates from a model archaeon, we identified AHA (AMPKγ-HPF from Archaea), a broadly conserved ribosome-associated protein factor composed of two distinct modules. Structural analyses showed that AHA’s C-terminal domain binds the small subunit, while its N-terminal region recognizes the large subunit, occluding the mRNA channel and the A- and P-tRNA binding sites and thereby enforcing translational silencing. Consistent with this proposed function, ΔAHA cells displayed reduced viability, depletion of ribosomal proteins during stationary phase, and impaired recovery upon return to growth. Phylogenetic analyses revealed that AHA’s C-terminal domain shares homology with the bacterial Hibernation Promoting Factor (HPF), indicating an origin in the last universal common ancestor (LUCA) and thereby identifying HPF as a universal hibernation module. Strikingly, we observed two AMP molecues bound to AHA’s N-terminal CBS-tetrad, which we found was structurally and evolutionary related to the eukaryotic energy sensor AMPKγ, thus linking energy sensing between archaea and eukaryotes. Together, these findings uncover a widespread archaeal ribosome hibernation factor and establish a direct evolutionary link between prokaryotic translational silencing and eukaryotic energy sensing.

## INTRODUCTION

Hibernation is a widely used strategy that allows organisms to survive environmental stresses, including starvation (Helena-Bueno *et al*., 2024a). Because protein synthesis is among the most energy-consuming processes in the cell, ribosome hibernation has emerged as a solution to reversibly halt translation without degrading ribosomal components, allowing rapid resumption of cell growth when conditions improve (Koli & Shetty, 2024; Njenga *et al*., 2023; Zegarra *et al*., 2023). In bacteria, hibernation is mediated by proteins such as hibernation-promoting factor (HPF) / RaiA (Agafonov *et al*., 1999), ribosome modulation factor (RMF) (Wada *et al*., 1990), and Balon (Helena-Bueno *et al*., 2024b), which block access to tRNA and mRNA binding sites or induce ribosome dimerization into translationally silent 100S particles (Beckert *et al*., 2017; Helena-Bueno *et al*., 2024b; Khusainov *et al*., 2017; Matzov *et al*., 2017; Polikanov *et al*., 2012). In eukaryotes, analogous factors, including LSO2 / CCDC124 (Wang *et al*., 2018), STM1 / SERBP1 (Ben-Shem *et al*., 2011), and MDF1 / MDF2 (Barandun *et al*., 2019), maintain ribosomes in an inactive but protected state during cellular stress.

Relative to bacteria and eukaryotes, archaeal ribosome hibernation (Hassan *et al*., 2025), and energy sensing and response remain poorly understood – alarmones analogous to bacterial (p)ppGpp have not been detected (van der Does *et al*., 2023), and the AMPK-dependent translation repression circuits like those conserved in eukaryotes have not been reported. Excitingly, recent work has begun to uncover proteins that bind ribosomes and appear to repress translation, such as the lineage-specific Dri protein from Thermoproteota (Nissley *et al*., 2025), the 30S-30S dimerization factor aRDF from Pyrococcus (Hassan *et al*., 2025), and additional hibernation factors related to that described here, which were reported in preprints released during the preparation of this manuscript (Madru *et al*., 2025; Nissley *et al*., 2026; Zhu *et al*., 2026).

In this work, we apply cryoEM to a crude lysate (May *et al*., 2025) prepared from cells entering stationary phase and identify a previously unrecognized ribosome-associated protein in the model haloarchaeon *Haloferax volcanii* (Pohlschroder *et al*., 2025). Structural analysis shows that this factor, which we term AHA (AMPKγ-HPF from Archaea) binds across the functional core of the ribosome and occludes the A- and P-tRNA binding sites, consistent with a role in translational silencing. Functionally, ΔAHA cells exhibited reduced viability upon exit from stationary phase and were profoundly impaired in their ability to compete with wild-type cells during repeated transitions between exponential growth and stationary phase. Remarkably, the protein is built from two distinct modules: a set of four cystathionine beta synthase (CBS) repeats (4xCBS) similar to the energy sensing γ-subunit of the eukaryotic AMPK complex, and a small-subunit–interacting domain homologous to bacterial HPF. We observe AMP molecules bound within the 4xCBS, in an arrangement similar to that observed in AMPKγ, suggesting that archaeal ribosome hibernation may be directly regulated by energy levels. The broad distribution of AHA-like 4xCBS-HPF factors we observed across archaea indicates that they were likely present in the last archaeal common ancestor. Moreover, the widespread occurrence of HPF-like domains in both archaea and bacteria implies that this hibernation module likely traces back to the last universal common ancestor (LUCA). The dual sequence and structural similarity of this archaeal factor to both HPF and AMPKγ suggests an evolutionary connection between ribosome hibernation and energy sensing, two fundamental strategies for surviving stress.

## RESULTS

### Discovery of a ribosome-associated protein in a model haloarchaeon

To investigate translation in the model haloarchaeon *Haloferax volcanii* (Pohlschroder *et al*., 2025), we applied cryoPRISM (May *et al*., 2025), a fast *ex vivo* cryoEM workflow that couples cell lysis, vitrification, data collection and analysis to determine high-resolution ribosomal structures in cell lysates, thereby preserving more native conditions than traditional cryoEM. We focused our analysis on cells transitioning from exponential to stationary phase growth, reasoning that this phase might reveal factors involved in modulating ribosome activity in response to nutrient limitation. Data collection and analysis of the resulting particles yielded a ∼2.5-Å resolution reconstruction of the *Hfx. volcanii* ribosome (**Figure S1**), the first high-resolution structure of a complete 70S ribosome from this model archaeal organism.

Because ribosomes are highly dynamic (Webster *et al*., 2023), consensus reconstructions inevitably represent a mixture of states (Sun *et al*., 2023). To address this heterogeneity, we performed 3D classification (**Figure S1**, see Methods), resulting in six major ribosome populations, each resolved at 2.4–2.8 Å resolution (**Table S1**) and distinguished by a varying constellation of factors and tRNAs in the intersubunit space. Four classes corresponded to canonical tRNA occupancy states: P/P-site only, A/A- and P/P-sites, P/P- and E/E-sites, and a hybrid A/P- and P-/E-configuration (**Figure 1A–D**). The remaining two contained an unexplained density, with one such state also bound by tRNA at the E-site (**Figure 1E–F**).

**Figure 1.**
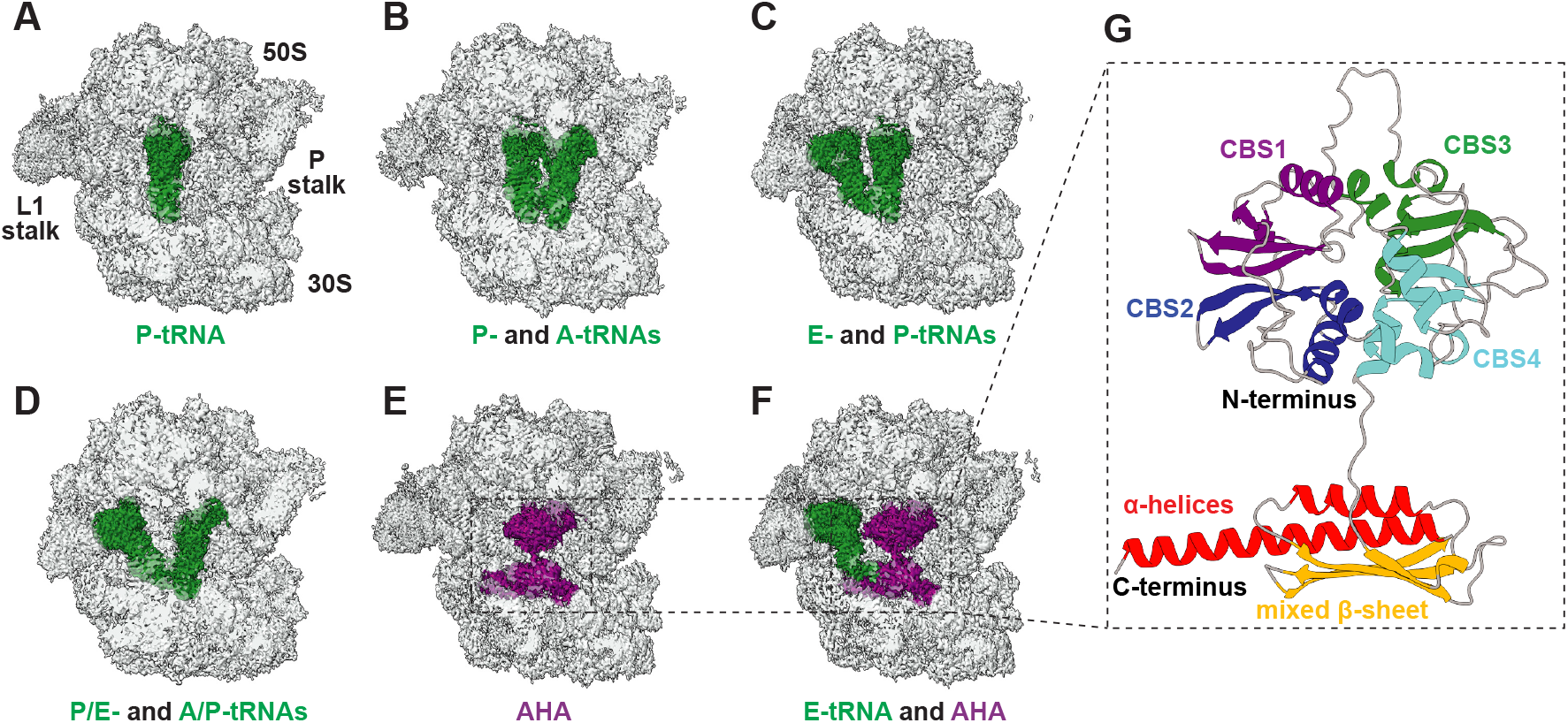
Discovery of a ribosome-associated protein in *Haloferax volcanii*. **(A-F)** CryoEM density maps of 70S ribosomes obtained via classification of ribosomal particles imaged directly from clarified *Hfx. volcanii* lysates. The 70S ribosome, tRNAs, and an additional unidentified density are shown in grey, green, and purple, respectively. Distinct tRNA occupancy states were observed, including (A) P/P-tRNA, (B) A/A- and P/P-tRNAs, (C) P/P- and E/E-tRNAs, (D) hybrid A/P- and P/E-tRNAs, and (E–F) ribosomes containing AHA, one of which also included an E-site tRNA (F). The locations of the 50S and 30S subunits and the L1 and P stalks are labeled in (A) for orientation. **(G)** Refined atomic model of the additional density unambiguously assigned to HVO_2384 by ModelAngelo (Jamali *et al*., 2024). The model, which is composed of an N-terminal domain with four consecutive CBS motifs and a C-terminal domain formed by a mixed β-sheet capped by two α-helices, is labeled colored by structural element. HVO_2384 N- and C-termini are indicated.

To identify this ribosome-bound factor, we performed automated model building with ModelAngelo (Jamali *et al*., 2024), providing this tool our high-resolution map and only rRNA and r-protein sequence information to facilitate an unbiased assignment of the density. Sequence analysis of the *ab initio* model unambiguously assigned the density to HVO_2384, a previously uncharacterized protein encoded in the *Hfx. volcanii* genome (**Figure S2**). This structure of HVO_2384 revealed two domains (**Figure 1G**): an N-terminal region containing four consecutive CBS motifs organized into two tandem Bateman modules (Bateman, 1997; Kemp, 2004) that packed against each other in a flower-like configuration, and a compact C-terminal fold consisting of a mixed parallel/antiparallel β-sheet capped by two α-helices. Our observation of HVO_2384 bound to ribosomes suggested that it may regulate translation.

### HPF is an ancient ribosome hibernation domain conserved across bacteria and archaea

To interrogate the functional and evolutionary relationships of HVO_2384, we performed remote homology and structure-based similarity searches using HHpred (Zimmermann *et al*., 2018) and Foldseek (van Kempen *et al*., 2024). HHpred enables sensitive profile–profile comparisons based on hidden Markov models (HMMs), thereby detecting distant evolutionary relationships at the sequence level (Söding, 2004), whereas Foldseek identifies structural homologs. These approaches converged on a consistent result: the N-terminal CBS tetrad of HVO_2384 exhibited significant sequence and structural similarity to CBS domain-containing regulatory proteins, most prominently the γ-subunit of eukaryotic AMPK-activated protein kinase (AMPKγ) (**Figure S3A**). These tools additionally uncovered homology between the C-terminal domain of HVO_2384 and bacterial hibernation-promoting factors (HPF) (**Figure S3B**). We thus concluded that HVO_2384 combines architectural features characteristic of metabolic CBS regulators with a C-terminal domain associated with ribosome hibernation factors. Given this dual similarity, bridging bacterial HPF and eukaryotic AMPKγ, we propose the name AHA (<u>A</u>MPKγ-<u>H</u>PF from <u>A</u>rchaea) for this *Hfx. volcanii* protein.

HPF and its paralog RaiA belong to a widely distributed and highly conserved bacterial protein family that mediates ribosome inactivation during stress (Agafonov *et al*., 1999; Chan *et al*., 2025; Helena-Bueno *et al*., 2024a). These factors bind to the small ribosomal subunit, where they occlude the A- and P-sites, thereby preventing tRNA and mRNA accommodation and effectively arresting translation (Polikanov *et al*., 2012). In many bacterial species, HPF homologs additionally promote the formation of translationally inactive 100S ribosome dimers (Beckert *et al*., 2017; Khusainov *et al*., 2017; Matzov *et al*., 2017), which protect ribosomes from degradation (Lipońska & Yap, 2021; Prossliner *et al*., 2021) and allow rapid reactivation once favorable conditions return. Consistent with the homology inferred above, the C-terminal domain of AHA adopted the same core secondary-structure elements characteristic of bacterial HPF, including the central β-sheet scaffold and flanking α-helices, although the terminal α-helix in AHA is longer than that in *E. coli* HPF (**Figure 2A**). To assess whether this structural similarity corresponded to a conserved mode of ribosome engagement, we superimposed bacterial ribosome-HPF complexes onto the *Hfx. volcanii* 70S-AHA structure, aligning the structures using only the core ribosomal RNA and proteins. Remarkably, the factors occupied an equivalent binding site within the small subunit (**Figure 2B**), partially filling the mRNA channel and occluding the A- and P-sites (**Figure 2C**). Our AHA-ribosome structure further revealed interactions between the HPF domain of AHA and 16S rRNA helices h18, h23, h24, h28, h30, h31, h34, and h44, as well as with ribosomal proteins uS7, uS11, and uS12 (**Figure 2D**). Importantly, the HPF domain of AHA and bacterial HPF bound to overlapping regions of the ribosome (**Figure 2D-E**) and used structurally equivalent residues to contact corresponding rRNA residues (**Figure S4**), indicating that archaeal and bacterial HPF domains recognize conserved ribosomal surfaces and likely inhibit translation through a shared steric-occlusion mechanism at critical tRNA binding sites.

**Figure 2.**
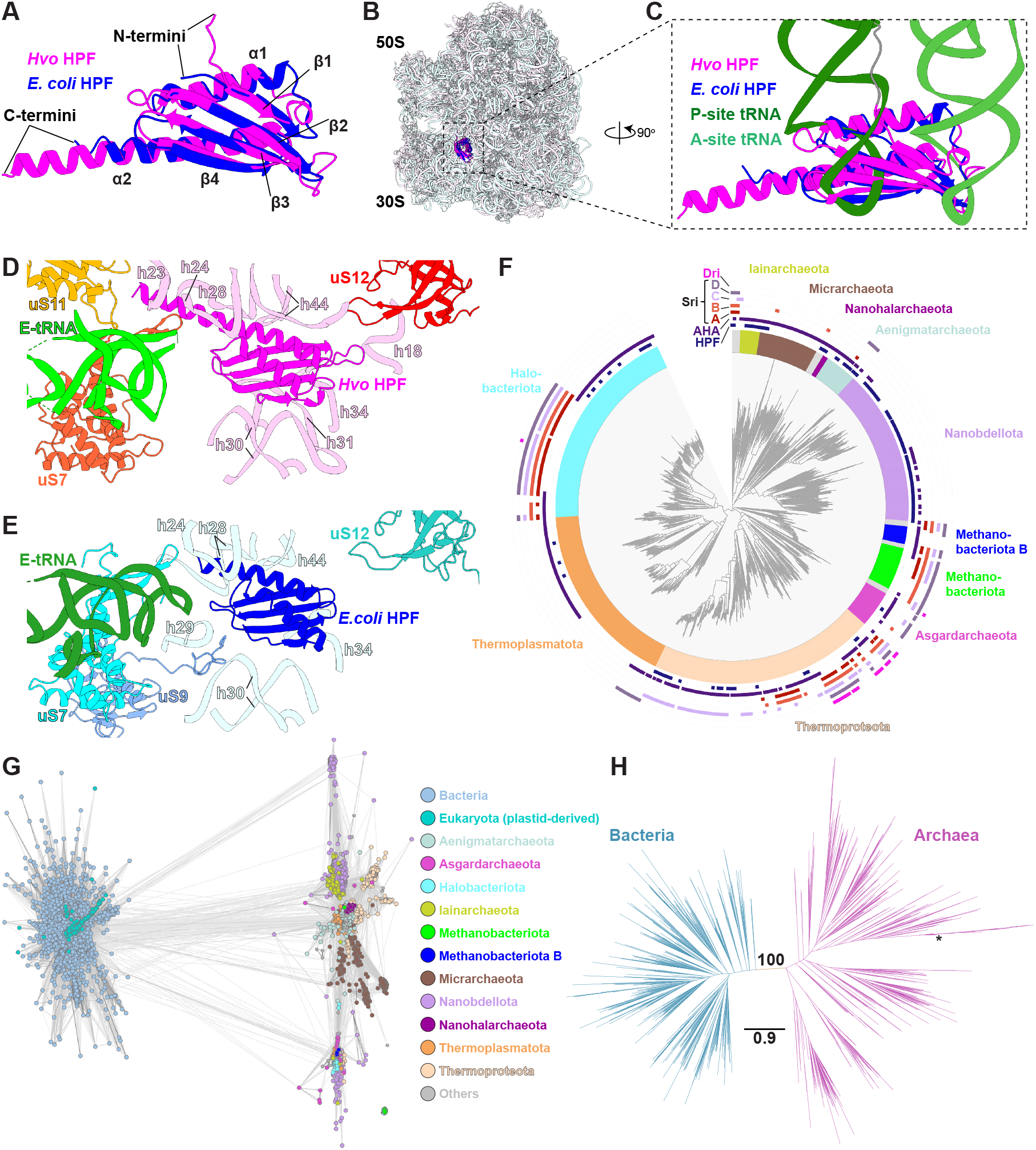
HPF is an ancient ribosome hibernation domain conserved across bacteria and archaea. **(A)** Structural superposition of the *Hfx. volcanii* (Hvo) AHA C-terminal HPF-like domain (pink) with *E. coli* HPF (blue; PDB: 6Y69). Secondary-structure elements and domain N- and C-termini are indicated. **(B-C)** Archaeal and bacterial HPF occupy the same site on the 30S subunit (archaeal and bacterial ribosomes in light pink and light blue, respectively), overlapping with the A- and P-tRNA binding sites (tRNAs colored in shades of green). **(D-E)** Comparison of the archaeal (D) and bacterial (E) HPF interactions with the 30S subunit. 16S rRNA helices and small-subunit proteins contacting the HPF domains are shown together with E-site tRNAs. A detailed comparison of the archaeal and bacterial HPF-30S interactions is provided in **Figure S4.** **(F)** Phylogenetic distribution across archaea of HPF- and CBS-containing proteins related to the domains found in AHA, mapped onto a species-level archaeal tree (center) derived from the Genome Taxonomy Database (Parks *et al*., 2021; Rinke *et al*., 2021). The thick inner ring denotes archaeal phyla, color-coded by taxonomic assignment. Concentric outer tracks indicate the presence of specific protein architectures, listed from inside to outside as: standalone HPF domains (no CBS fusion), AHA-type proteins (4xCBS-HPF fusion), SriA-SriD proteins (4xCBS without HPF) (Nissley *et al*., 2026), and Dri proteins (two consecutive 4xCBS modules) (Nissley *et al*., 2025). Tracks, colored by the key, mark genomes encoding the indicated architectures. **(G)** CLANS cluster-map analysis of HPF homologues based on all-against-all pairwise sequence similarities. Each dot represents one protein, colored by taxonomic group (bacteria – light blue; eukaryotes - aqua; archaea – colored by phylum). Line intensity reflects sequence similarity, with darker lines indicating higher similarity (*i*.*e*., lower BLAST p-values). HPF homologs from bacteria (left cluster) and archaea (right cluster) clearly separate, and the archaeal HPF domains largely segregate according to their phylum-level assignments. **(H)** Maximum-likelihood phylogenetic tree of the HPF superfamily across bacteria (blue) and archaea (pink), with 1,000 HPF homologues sampled across domains analyzed. The scale bar indicates amino-acid substitutions per site. The central branch separating archaeal and bacterial sequences is maximally supported, with an ultrafast bootstrap value of 100 (shown) and SH-aLRT support of 99.7. An asterisk indicates the *Hfx. volcanii* AHA HPF sequence.

To investigate the phyletic distribution of AHA and the evolutionary origins of HPF-like ribosome hibernation domains in archaea, we surveyed 6,968 archaeal genomes spanning 21 phyla, using the HPF domain of *Hfx. volcanii* AHA as the primary query to specifically track ribosome-associated homologs (see Methods). In total, we identified 3,160 AHA homologs distributed across 2,723 distinct archaeal species spanning 17 phyla, and 926 canonical HPF homologs (*i*.*e*., proteins with standalone HPF domains lacking the N-terminal CBS tetrad) present in 814 species across 12 phyla. Multiple AHA or HPF paralogs were often observed within individual genomes (**Figure 2F**). Notably, AHA and canonical HPF homologs were also detected within Asgardarchaeota, including Hodarchaeales, an Asgard lineage frequently discussed in the context of eukaryogenesis (Eme *et al*., 2023).

HPF-containing proteins were detected across multiple major archaeal phyla (**Figure 2F**). We considered that this broad distribution could reflect three plausible evolutionary scenarios: (*i*) an ancient origin predating the split of bacteria and archaea (*i*.*e*., presence in the LUCA); (*ii*) a single early acquisition from bacteria near the archaeal root; or (*iii*) repeated horizontal gene transfer (HGT) events from bacteria into different archaeal clades. To distinguish among these possibilities, we performed sequence clustering and maximum-likelihood phylogenetic analyses of archaeal and bacterial HPF homologs.

CLANS clustering (Frickey & Lupas, 2004; Zimmermann *et al*., 2018) of HPF homologs, based on all-against-all pairwise sequence similarity, separated HPF sequences into two domain-level clusters (**Figure 2G**). One dense cluster contained bacterial HPFs and plastid-derived eukaryotic homologs, consistent with acquisition of HPF by eukaryotes via the cyanobacterial endosymbiont (Sharma *et al*., 2010; Swift *et al*., 2020; Tanaka *et al*., 2025). The other cluster comprised archaeal HPFs and exhibited clear phylum-level substructure. Across the dataset, archaeal sequences showed their strongest similarities to other archaeal homologs rather than to bacterial HPFs, a pattern incompatible with recent or lineage-restricted bacterial-to- archaeal HGT.

To test this separation more explicitly, we inferred a maximum-likelihood tree from 1,000 HPF homologs sampled in a taxonomy-balanced manner across archaea and bacteria (**Figure 2H**). The unrooted tree recovered two deeply divergent, domain-specific clades corresponding to archaeal and bacterial sequences, with strong statistical support (ultrafast bootstrap = 100; SH-aLRT = 99.7). Each clade was cohesive, and we observed no cross-domain intermixing. Archaeal HPF homologs also displayed greater internal sequence divergence than bacterial homologs, consistent with long-term independent evolution within each domain rather than recent, lineage-restricted transfer.

Together, these observations indicated that HPF was already present in the last archaeal common ancestor (LACA). Because HPF is also broadly conserved across bacteria (Chan *et al*., 2025) and inferred to have been present in the last bacterial common ancestor (LBCA), the most parsimonious explanation is that HPF originated in, or was inherited from, the LUCA. Accordingly, we posit that AHA represents the archaeal branch of an ancient ribosome hibernation factor family conserved across bacteria and archaea.

### AHA is required for ribosome hibernation and survival in stationary phase

Given that bacterial HPF binds the ribosomal core and promotes hibernation by arresting translation as cells enter stationary phase, we hypothesized that AHA may play a similar role in archaea. To test this hypothesis, we generated an *Hfx. volcanii* knockout of the HVO_2384 gene encoding AHA. When cultures were inoculated from cells grown in exponential phase, wild-type and ΔAHA strains grew similarly (**Figure 3A**). In contrast, after inoculation with cells in stationary phase, ΔAHA cells exhibited a marked delay in resuming logarithmic growth compared to wild-type cells (**Figure 3B**). Complementation with the HVO_2384 gene rescued this phenotype (**Figure 3B**), confirming that the defect was specific to loss of AHA.

**Figure 3.**
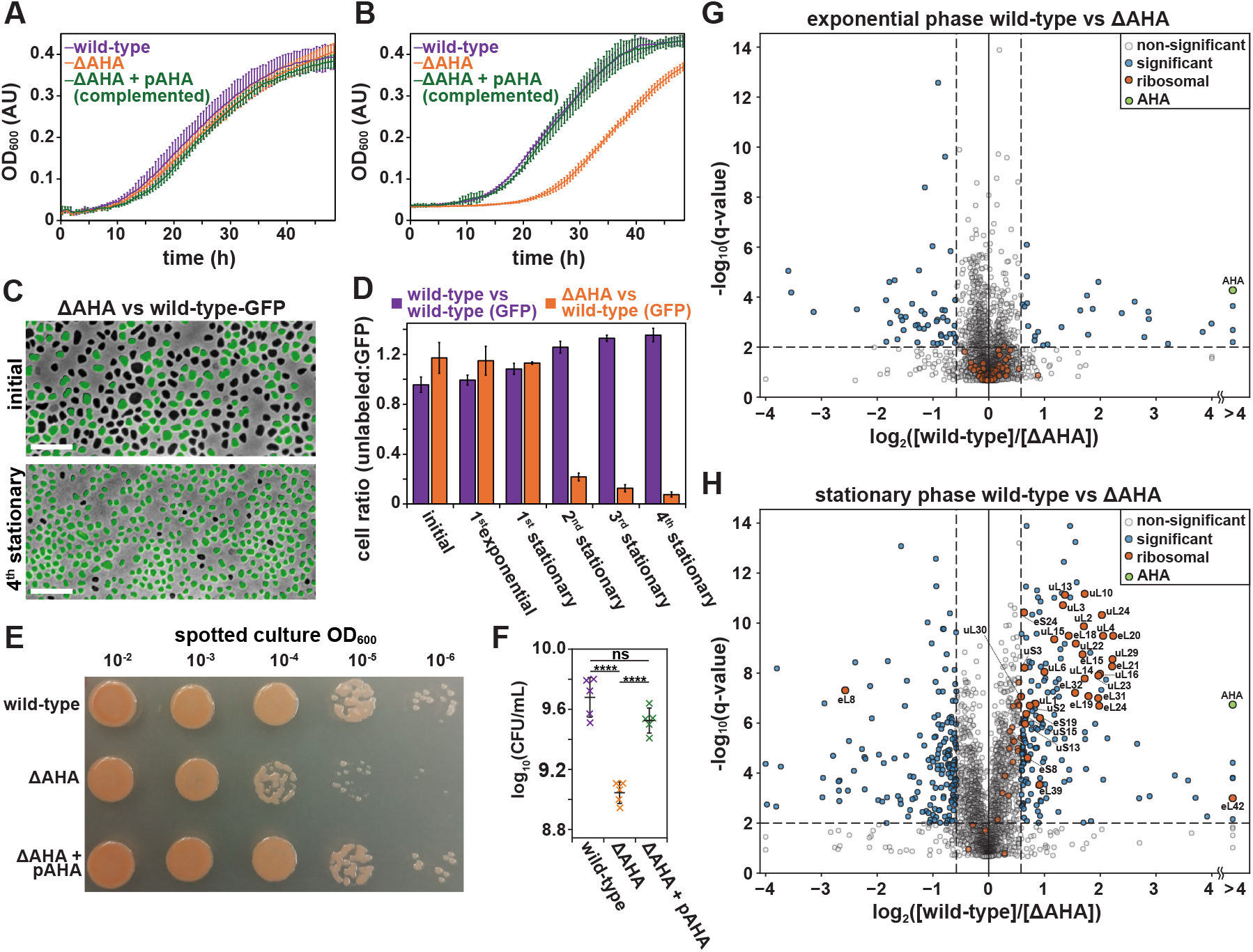
AHA is required for survival in stationary phase. **(A-B)** Growth, as measured by monitoring light scattering (OD_600_) of wild-type (purple), ΔAHA (orange), and complemented ΔAHA + pAHA (green). Assays were initiated by diluting exponentially growing cultures (OD_600_ ≈ 0.4) into Hv-Cab media to a final OD_600_ = 0.05, (A) or by such inoculation from stationary-phase cultures isolated at OD_600_ ≈ 4.0 (B). Error bars mark the standard deviation of biological replicates (n ≥ 3). **(C)** Competitive growth assays between a wild-type strain constitutively expressing GFP and either a wild-type or a ΔAHA strain, with competing strains mixed in equal ratio. Overlay of phase-contrast and GFP fluorescence images showing ΔAHA (non-fluorescent) and wild-type-GFP (fluorescent) cells immediately after mixing (top, “initial”) and after four consecutive growth–dilution cycles to stationary phase (bottom, “4^th^ stationary”). Scale bars, 10 µm. **(D)** Ratio of non-fluorescent (wild-type or ΔAHA) to fluorescent (wild-type-GFP) cells across growth successive cycles as described in (C). Images were collected immediately after mixing (initial), late in the first round of logarithmic growth (1^st^ exponential) and subsequently at various (1^st^-4^th^) rounds of stationary phase. Bars mark the mean ± standard deviation of three independent competitions. **(E)** Colonies of wild-type, ΔAHA, and complemented ΔAHA + pAHA strains plated from stationary-phase cultures. Note that under these conditions, ΔAHA colonies were heterogeneous in size and generally smaller than those of wild-type or complemented strains. **(F)** Viable colony counts (CFU·mL^−1^, log_10_ scale) of the enumerated strains from stationary-phase cultures. Each cross represents one biological replicate (n = 5); mean ± SD shown with black ticks. Statistical comparisons were performed using a two-tailed Welch’s t-test on log_10_(CFU·mL^−1^) values; **** notes p < 1×10^−4^; ns notes not significant (p ≥ 0.05). **(G-H)** Volcano plot highlighting changes in protein levels between wild-type and ΔAHA *Hfx. volcanii* grown in exponential (G) or stationary (H) phase. The log_2_-fold change of protein abundance (x-axis) is plotted against the statistical significance (y-axis) across 3-4 replicates for each condition (see Methods). Blue points represent “significant” proteins exhibiting a greater than 1.5-fold change in abundance (x-axis) and high statistical significance (q-value ≤ 0.01, y-axis). Ribosomal proteins are shown in orange, with larger circles indicating those that pass the significance criteria. AHA is colored green.

We next asked whether AHA deficiency conferred a competitive disadvantage relative to wild-type cells. Specifically, ΔAHA cells were co-cultured with the parental strain expressing cytoplasmic msfGFP (Pédelacq *et al*., 2006) under a constitutive promoter (wild-type-GFP; **Figure 3C**). In parallel, wild-type-GFP cells were co-cultured with the unlabeled parental strain as a control assay. Over four cycles of growth to stationary phase followed by dilution into fresh media, wild-type-GFP cells outcompeted ΔAHA, whose relative abundance steadily declined at a rate corresponding to a selection coefficient s = −0.86, or a ≈58% reduction per stationary-phase cycle (**Figure 3D**).

To clarify whether the regrowth defect of ΔAHA reflected loss of viability or delayed recovery from stationary phase, we quantified the number of colony-forming units after cells had entered stationary phase. ΔAHA cultures produced ≈4.4-fold fewer colonies than wild-type, and ΔAHA colonies were generally smaller in size (**Figure 3E-F**), indicating that the loss of AHA compromised cell survival during stationary phase and, additionally, impaired subsequent recovery and growth.

To investigate the molecular basis of the ΔAHA deficiency in exiting stationary phase, we compared the proteomes of wild-type and ΔAHA cells in exponential and stationary phase using quantitative mass spectrometry (**Figure 3G–H**, see Methods). As expected, AHA was undetectable in the knockout strain (**Figure S5** and **Source Data Table 1**), confirming the specificity of the deletion. In contrast to classic bacterial hibernation factors whose expression levels are strongly growth-phase dependent (Helena-Bueno *et al*., 2024b; Izutsu *et al*., 2001b; Maki *et al*., 2000; Yamagishi *et al*., 1993), AHA levels were largely unchanged in wild-type cells grown in either exponential or stationary phase, suggesting an alternative mechanism of regulation (**Figure S5**). Moreover, relatively few changes were detected across the proteome in cells growing exponentially in rich media (**Figure 3G**), indicating that AHA exerted a restricted influence on the expressed proteome under these conditions. Notably, however, in stationary phase, ΔAHA cells exhibited widespread remodeling of their proteome relative to wild-type cells, including significant changes in pathways related to DNA replication and repair, phosphate, polyphosphate and amino acid metabolism, cell envelope biogenesis and transport, and redox and general stress responses (**Source Data Table 1**). Most strikingly, ΔAHA cells in stationary showed a coordinated depletion of 32 ribosomal proteins, predominantly from the large subunit (**Figure 3H**). This pattern was strongly indicative of ribosome destabilization and degradation in the absence of AHA, consistent with the view that AHA protects ribosomes during hibernation and prevents their loss under energy-limiting conditions.

### The eukaryotic energy sensing protein AMPKγ evolved from archaeal factors related to AHA

The N-terminus of AHA contains four consecutive CBS motifs that formed a compact domain observed in complex with the ribosomal large subunit (**Figure 4A**), spanning the A- and P-tRNA sites (**Figure 4B**). We observed contacts between the 4×CBS region and 23S rRNA helices H69, H70, H71, H73-H74 loop, H74, H80/P-loop, H89, H90, H92/A-loop, and H93, the 50S protein uL16 and the 30S subunit uS19 (**Figure 4C**). Additionally, the third CBS motif contained an extended loop 3 that was inserted into the peptidyl transferase center (PTC; **Figure 4D**), where it interacted with universally conserved nucleotides that line the PTC active site (G2061/G2094, U2506/U2533, G2583/G2610, U2584/U2611, U2585/U2612 and A2602/ A2629; *E. coli* and *Hfx. volcanii* numbering, respectively), and which play key roles in positioning the A- and P-site substrates during peptide-bond formation (Polacek & Mankin, 2005; Voorhees *et al*., 2009). Interestingly, A2602/ A2629, which aids in triggering peptidyl-tRNA hydrolysis during translation termination (Amort *et al*., 2007) was inserted into the cavity created by loop 3 (**Figure 4D**, bottom). When compared to the induced and uninduced PTC conformations described by Schmeing *et al*., 2005, the AHA-bound ribosome adopted a non-canonical “partially induced” state: U2585/U2612 remained uninduced-like; U2506/U2533 and U2584/U2611 were only partially shifted toward the induced configuration; and A2602/A2629 was repositioned into an alternative orientation distinct from canonical elongation or termination geometry.

**Figure 4.**
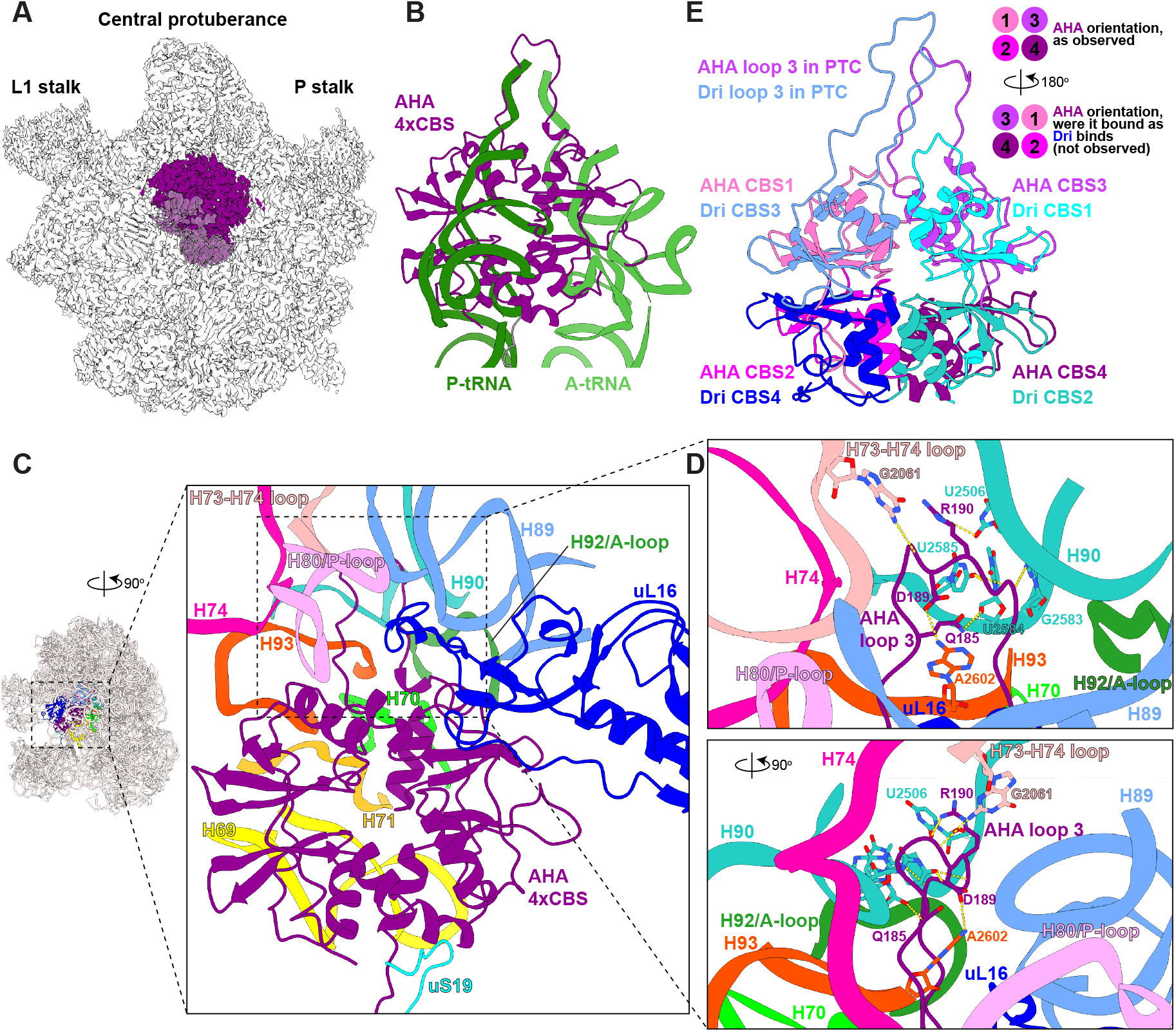
The AHA N-terminal 4xCBS domain binds the large subunit and inserts into the peptidyl transferase center. **(A)** CryoEM reconstruction of the *Hfx. volcanii* 50S subunit (grey; 30S omitted for clarity) bound to the AHA N-terminal 4×CBS domain (purple). The locations of the central protuberance and L1 and P stalks are labeled for orientation. **(B)** Structural overlap of the AHA 4×CBS domain (purple) with A- and P-site tRNAs (green shades), illustrating that AHA occupies these canonical tRNA-binding regions of the large subunit. **(C)** The 4×CBS domain forms extensive contacts with multiple 23S rRNA helices (H69, H70, H71, H73-H74 loop, H74, H80/P-loop, H89, H90, H92/A-loop and H93) and engages neighboring proteins uS19 and uL16. **(D)** A long loop within the third CBS motif (“loop 3”) penetrates deeply into the peptidyl transferase center (PTC), where it interacts with PTC universally conserved nucleotides (G2061, U2506, G2583, U2584, U2585 and A2602; *E. coli* numbering), which position the A- and P-site substrates during pep-tide-bond formation. Several AHA side chains contacting these nucleotides are shown, and interactions are indicated by yellow dashed lines. Notably, A2602 inserts into the cavity created by loop 3 (bottom panel). **(E)** Comparison of the AHA 4×CBS domain (purple/pink) with the archaeal hibernation factor Dri from Thermoproteota (blue; PDB: 9E6Q). Both proteins insert an equivalent loop into the PTC, but their CBS modules bind the large subunit in opposite orientations, necessitating a ∼180° rotation to align their CBS motifs, as depicted in the inset.

Notably, the AHA CBS region in our structure resembled the N-terminal domain of the recently described Thermoproteota-specific hibernation factor Dri, which also contains four CBS repeats and binds at the A- and P-tRNA sites (Nissley *et al*., 2025). However, despite this similarity, AHA and Dri bound to the ribosomal large subunit in different orientations, with a ∼180° rotation required to align their equivalent CBS motifs (**Figure 4E**).

As noted, Foldseek- and HHpred-based analyses of the AHA CBS region revealed high-confidence structural similarity to the eukaryotic AMPKγ, and such similarities were also apparent when aligning the structures (**Figure 5A**). Given the AHA - AMPKγ homology, and AMPKγ’s well defined role in nucleotide sensing (Smiles *et al*., 2024; Steinberg & Hardie, 2023), we next examined whether AHA might also bind nucleotides. Direct inspection of our cryoEM maps of AHA•ribosome complexes revealed density (**Figure 5B**, right) consistent with two AMP molecules bound at the junction of CBS3 and CBS4 (**Figure 5C**). Strikingly, these AMP-binding pockets directly corresponded to the canonical regulatory sites of AMPKγ (**Figure 5B**, left) (Chen *et al*., 2012; Gu *et al*., 2017; Hawley *et al*., 2024; Yan *et al*., 2021), with each protein recognizing nucleotides through a combination of conserved and lineage-specific residues (**Figure 5C, S6**).

**Figure 5.**
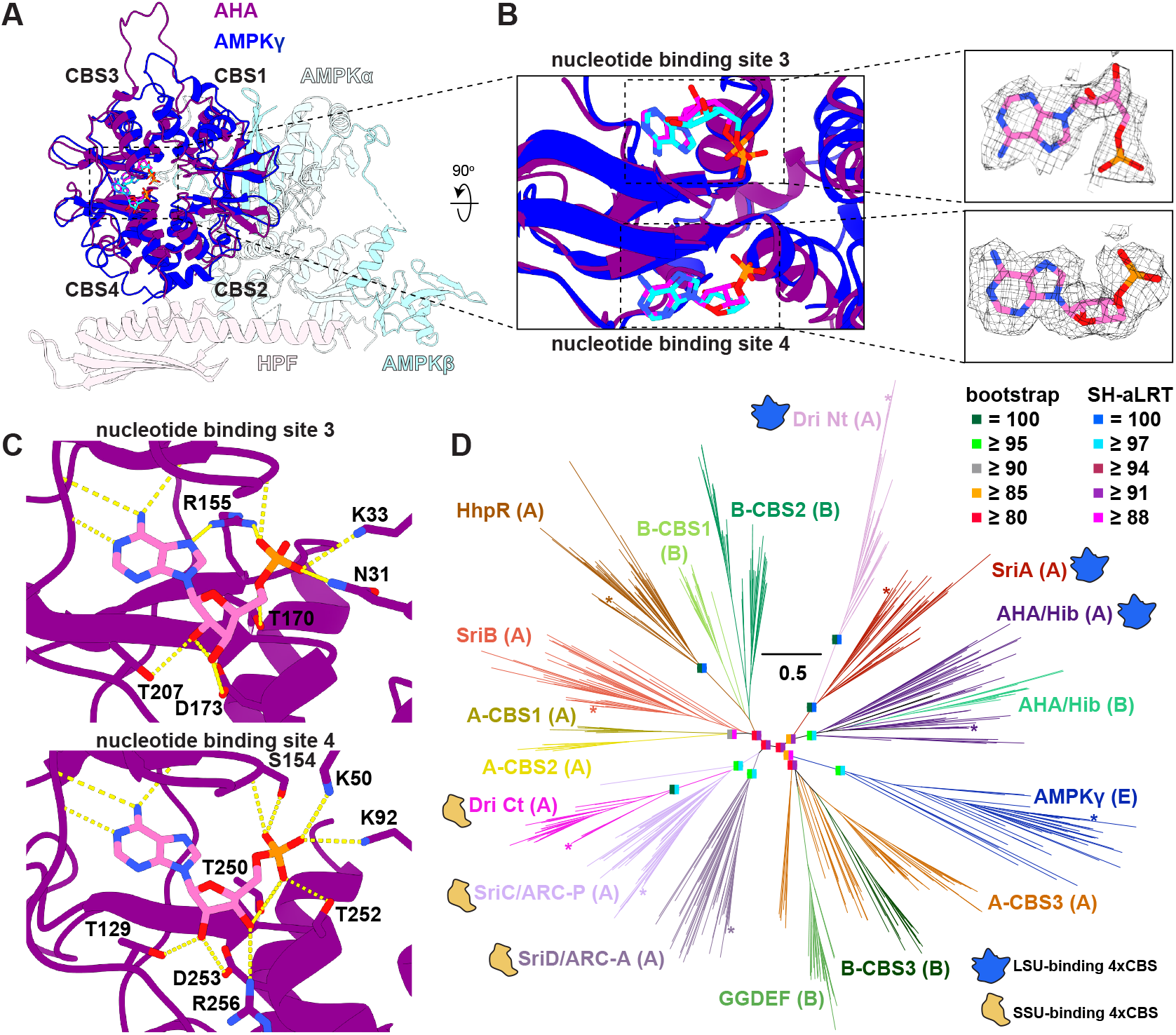
The AHA N-terminal 4×CBS domain binds AMP and closely resembles eukaryotic AMPKγ. **(A)** Structural superposition of the AHA 4×CBS domain (purple) with the eukaryotic AMPKγ subunit (dark blue). The remaining components of the AMPK complex, AMPKα and AMPKβ, are shown in light blue shades (PDB: 6B1U). AHA’s C-terminal HPF domain is displayed in light pink for reference. **(B)** Two AMP molecules (carbon atoms in pink) are bound between CBS3 and CBS4 of AHA, occupying positions equivalent to nucleotide-binding sites 3 and 4 of AMPKγ, whose bound AMP carbons are shown in cyan (left). Insets show the corresponding cryoEM densities for the two AHA’s AMP molecules (right). Oxygen, nitrogen, and phosphorus atoms are colored red, blue, and orange, respectively. **(C)** Detailed views of the AMP-binding pockets in AHA. Protein residues forming hydrogen bonds or electrostatic interactions with the nucleotides are labeled, and interactions are indicated by yellow dashed lines. Most purine-ring contacts are mediated by backbone atoms (not shown for clarity). **(D)** Unrooted maximum-likelihood phylogenetic tree of the CBS-domain protein superfamily related to the N-terminal region of AHA. The scale bar represents amino-acid substitutions per site. Ultrafast bootstrap and SH-aLRT support values are shown for selected internal nodes, particularly those defining major protein families (see colored rectangles). Asterisks mark experimentally characterized reference proteins (species; UniProt accession): the eukaryotic energy sensor AMPKγ (*Homo sapiens*; P54619); the archaeal transcriptional regulator HhpR (*Pyrococcus yayanosii*; F8AFU3); and archaeal ribosome hibernation factors, including AHA (*Hfx. volcanii*; D4GWN2), SriA–D (*Methanosarcina acetivorans*; Q8TH74, Q8TH73, Q8TH72, Q8TH71), and Dri (N- and C-terminal domains analyzed separately for this phylogeny; *Pyrobaculum calidifontis*; A3MX58). Asterisk colors correspond to their respective family labels. The tree resolves several previously uncharacterized CBS-domain subfamilies, including archaeal CBS1–3 (A-CBS1–3), bacterial CBS1–3 (B-CBS1–3), and bacterial 4×CBS–GGDEF fusion proteins (GGDEF). Although AHA is primarily archaeal, a limited number of bacterial homologues are also observed. Letters in parentheses indicate the domain of life in which each family is predominantly found: (A) Archaea, (B) Bacteria, and (E) Eukaryota. Proteins with experimentally validated binding of the 4×CBS domain to the ribosomal large (blue) or small (yellow) subunit are indicated.

To explore the evolutionary relationship between archaeal ribosome hibernation and eukaryotic energy sensing, we investigated the evolutionary origins of the CBS tetrad of AHA and asked whether AMPKγ may have emerged from an archaeal CBS-tetrad lineage. This analysis was prompted by reciprocal HHpred searches, in which AHA and AMPKγ consistently ranked among each other’s strongest matches, suggesting a specific relationship beyond generic similarity among CBS-containing proteins. To define the broader sequence landscape of AMPKγ-like CBS tetrads, we used an AMPKγ profile HMM to survey archaeal, bacterial, and eukaryotic databases. The retrieved homologs were then curated into a non-redundant dataset and analyzed using CLANS clustering based on all-against-all pairwise sequence similarity (see Methods). The resulting map resolved two clearly separated regions (**Figure S7**). One region comprised eukaryotic AMPKγ and related homologs, forming a compact and cohesive cluster. The second region consisted of a central “core” dominated by archaeal CBS-tetrad proteins, from which multiple subclusters radiated.

This archaeal-centered core included the AHA CBS tetrad together with the recently described ribosome hibernation factors SriA–D (Nissley *et al*., 2026) / ARC-AP (Zhu *et al*., 2026), which bear a CBS tetrad; Dri (Nissley *et al*., 2025), which contains two CBS tetrads; and the *Pyrococcus yayanosii* transcription factor HhpR (Li *et al*., 2025), which is composed of one CBS tetrad (**Figure S8**). Notably, the AHA subcluster also contained a small number of bacterial AHA-like sequences, primarily from Patescibacteria (CPR) lineages, which were embedded within the archaeal cluster rather than forming a distinct bacterial branch. The broader central region additionally contained other CBS-tetrad families, including bacterial GGDEF (Jia *et al*., 2025) -associated CBS proteins and several smaller archaeal and bacterial groups (A- and B-CBS1–3). These formed distinct subclusters separable from the Sri/AHA lineage, indicating that the CBS-tetrad architecture has diversified into multiple functional contexts, and is not restricted to ribosome regulation.

Mapping these CBS subfamilies onto the archaeal species tree (**Figure 2F**) revealed contrasting distribution patterns. Dri showed a restricted phylogenetic occurrence, consistent with previous reports (Nissley *et al*., 2025), whereas AHA displayed a broad and phylogenetically widespread distribution across archaeal superphyla. In contrast, SriA–D exhibited a more lineage-structured distribution. Strikingly, AHA and Sri homologs show largely non-overlapping phylogenetic patterns, suggesting partial functional redundancy and possible evolutionary replacement or competition between these hibernation systems containing CBS-tetrads. Overall, 5,080 of the 6,968 archaeal species in our GTDB dataset encode at least one of these ribosome hibernation modules (AHA, SriA–D, Dri, or canonical HPF). Collectively, these factors are represented across all 21 archaeal phyla sampled, indicating that such ribosome hibernation systems are nearly ubiquitous at the phylum level, despite strong lineage-specific partitioning at finer taxonomic scales.

To resolve relationships among these CBS-containing families, we performed maximum-likelihood phylogenetic analyses (**Figure 5D**, see Methods). The resulting tree revealed several notable features. First, ribosome-associated CBS proteins segregated into two major functional clusters: (*1*) factors reported to act on the small subunit (*i*.*e*., SriC, SriD, ARC-A/P, and the Dri C-terminal domain), and (*2*) those associated with the large subunit (*i*.*e*., AHA, SriA, and the Dri N-terminal domain). Second, Dri appeared to have originated from a fusion event involving SriA-like and SriC/ARC-P–like ancestors, consistent with its two-CBS-tetrad architecture. Third, according to this tree, AHA likely evolved from a SriA-like ancestor that subsequently fused to an HPF domain, giving rise to the dual-module hibernation factor described here. Fourth, several bacterial CBS-tetrad families branched within archaeal clades and displayed either lineage-restricted distributions or phylogenetically discontinuous patterns across bacteria, features more consistent with horizontal acquisition than with deep vertical bacterial ancestry. Notably, several of these bacterial families possess the extended loop characteristic of AHA CBS3 (**Figure S8**), raising the possibility that they may likewise be ribosome-associated.

Our CBS-tetrad phylogeny consistently placed eukaryotic AMPKγ within a clade comprising an uncharacterized archaeal family (A-CBS3), together with a limited number of bacterial representatives. Although statistical support for some deep branches was moderate and the precise placement of AMPKγ was not fully resolved, the overall topology is nevertheless compatible with an archaeal origin of the AMPKγ lineage. In this framework, the CBS tetrad of AHA can be interpreted as deriving from a broader archaeal CBS-tetrad radiation that also gave rise to the eukaryotic AMPKγ family. The limited and uneven bacterial representation is most parsimoniously explained by horizontal transfer rather than by an ancient bacterial diversification. More broadly, these data support a model in which multiple CBS-tetrad ribosome-associated factors co-existed early in archaeal evolution, followed by lineage-specific duplications, fusions, and differential retention, ultimately linking archaeal ribosome hibernation systems to the emergence of eukaryotic energy sensing.

## DISCUSSION

Ribosome hibernation is a broadly conserved strategy that allows cells to survive periods of nutrient limitation by reversibly restricting translation (Koli & Shetty, 2024; Smith *et al*., 2022; Usachev *et al*., 2020). Whereas bacterial and eukaryotic hibernation factors have been extensively characterized (Helena-Bueno *et al*., 2024a; Koli & Shetty, 2024; Smith *et al*., 2022), archaeal counterparts remain poorly understood, with the exceptions of the lineage-specific Dri (Nissley *et al*., 2025), aRDF (Hassan *et al*., 2025), and Hib/SriA-D/ARC-A-P, reported recently in preprints (Madru *et al*., 2025; Nissley *et al*., 2026; Zhu *et al*., 2026). By applying cryoPRISM (May *et al*., 2025) directly to *Hfx. volcanii* lysates harvested from cells entering stationary phase, we identified AHA, a ribosome-associated protein that binds at the functional core of the ribosome. Notably, this work illustrates the utility of near-native cryoEM, and workflows such as cryoPRISM in particular, to uncover previously uncharacterized proteins associated with core macromolecular complexes, including regulatory factors that are often lost during purification. Moreover, it highlights the ability of structural methods in generating testable hypotheses, such as the role of AHA in supporting ribosome hibernation, which we directly tested here.

The broad phylogenetic distribution of HPF domains we observed across archaeal superphyla indicates that ribosome hibernation is not a recent or lineage-specific adaptation, but rather a deeply conserved feature of archaeal biology, likely present in the last archaeal common ancestor. Moreover, since HPF is also ubiquitous and highly conserved in bacteria (Chan *et al*., 2025), the most parsimonious explanation is that this factor originated in the LUCA. HPF therefore represents the first universal ribosome hibernation factor described across both domains of life, bacteria and archaea (Nobs *et al*., 2022; Williams *et al*., 2020).

Consistent with this evolutionary conservation, we found that the absence of AHA in *Hfx. volcani*i led to pronounced regrowth defects after stationary phase, reduced viability, depletion of ribosomal proteins, and decreased competitive fitness, phenotypes that closely mirror those associated with HPF deficiency in bacteria. Furthermore, we observed both HPF and AHA bound across the A- and P-sites, occluding the mRNA channel, and extending into the E-site. These interactions are fundamentally incompatible with translation, consistent with the notion that AHA, like bacterial HPF, functions as a direct translational inhibitor that stabilizes ribosomes in a hibernating state during energy limitation.

An important distinction, however, emerges when considering how AHA activity may be regulated. In bacteria, the induction of ribosome hibernation is tightly coupled to transcriptional control of the abundance of hibernation factors through the stringent response and the alarmone (p)ppGpp (Eymann *et al*., 2002; Izutsu *et al*., 2001a; Nagarajan *et al*., 2025). This regulatory system is absent in archaea (van der Does *et al*., 2023). Notably, we found that AHA was robustly expressed at similar levels in both exponential and stationary growth, suggesting that archaeal ribosome hibernation is not governed primarily by transcriptional regulation. Instead, AHA carries a distinctive evolutionary signature that points to a different regulatory logic: its N-terminal four-CBS domain closely resembles the γ-subunit of the eukaryotic AMPK, and we observed AMP bound at AHA’s conserved pockets that correspond to the principal regulatory sites of AMPKγ.

This observation raises the possibility that the AHA·ribosome interaction is regulated by adenine nucleotide binding. Indeed, the relative levels of ATP, ADP, and AMP are canonical readouts of cellular energy status (Steinberg & Hardie, 2023), and translational downregulation is expected to be advantageous under low-energy conditions. We therefore hypothesize that adenine nucleotide binding modulates the affinity of AHA for the ribosome and, consistent with this hypothesis, a recent preprint reported that ARC-A/P proteins are released from ribosomes in the presence of ATP (Zhu *et al*., 2026). How then might AHA couple adenine nucleotide binding to differential affinity for its ribosome binding site? The proximity of AHA’s nucleotide-binding sites to loop 3, an extended element that inserts into the peptidyl transferase center, suggests a biochemical mechanism that links these binding events: AMP binding stabilizes and exposes loop 3, thereby increasing AHA’s affinity for the ribosome to promote ribosome hibernation under energy stress.

Taken together, our work suggests that CBS domains originally contributed to ribosome-level energy control and were later co-opted into the eukaryotic AMPK complex to regulate energy metabolism on a cellular scale. In this view, AHA sits at an evolutionary crossroads: its C-terminal HPF-like domain exemplifies an ancient, LUCA-era strategy for stabilizing inactive ribosomes, whereas its N-terminal CBS tetrad anticipates the architecture of eukaryotic AMPKγ. By uniting these two modules on a single archaeal factor, AHA provides a direct evolutionary and mechanistic link between prokaryotic ribosome hibernation and the eukaryotic energy-sensing machinery. This framework suggests that energy-responsive translational silencing is not a recent innovation but a deeply rooted principle that was later expanded and rewired as regulatory networks became more complex.

Looking forward, several questions follow directly from this work. A central priority will be to define quantitatively how adenine nucleotides modulate AHA·ribosome binding and whether changes in adenine nucleotide ratios are sufficient to switch AHA into a ribosome-hibernation-promoting state *in vivo*. It will also be important to determine whether additional archaeal CBS-tetrad proteins we identified bioinformatically act as ribosome-associated regulators, how their activities intersect with hibernation systems such as Dri, aRDF, SriA-D/ARC-AP and AHA/Hib, or whether they serve other roles in maintaining cellular homeostasis (e.g., in regulating transcription). In this light, our structural and functional characterization of AHA as a widespread archaeal ribosome hibernation factor supports the idea that transient ribosome inactivation is an ancient strategy for conserving energy, which our phylogenetic analyses indicate was already established in the earliest common ancestors of contemporary life.

## MATERIALS AND METHODS

### *Haloferax volcanii* cell growth and strain generation

*Hfx. volcanii* strains were grown in semi-defined Hv-Cab medium (de Silva *et al*., 2021), supplemented with 50 μM uracil when required. Cultures were incubated at 42 °C for liquid growth and 45 °C for growth on solid medium. Oligonucleotides, plasmids, and strains used in this study are listed in Table S2. Plasmids were assembled by isothermal Gibson cloning (Gibson *et al*., 2009), transformed into chemically competent *E. coli* DH5α, and selected on LB plates containing 100 μg/mL carbenicillin. Purified plasmids were subsequently introduced into *Hfx. volcanii* using the polyethylene glycol (PEG)-mediated transformation method described by (Bitan-Banin *et al*., 2003). The ΔAHA gene knockout was generated using the classical “pop-in, pop-out” recombination strategy (Bitan-Banin *et al*., 2003) in the parental H26 (ΔpyrE2) strain. Counter-selection for loss of the integrated plasmid was performed on medium containing 5-fluoroorotic acid (5-FOA). An AHA complementation plasmid was generated by cloning the AHA gene into the replicative pAL750 vector series (Rados *et al*., 2023). This plasmid, together with the empty pAL750 vector, was transformed into the H26 and ΔAHA backgrounds as described above, yielding the strains H26 pAL750 (aBL639), ΔAHA pAL750 (aBL640), and ΔAHA pAL750-AHA (aBL641), hereafter referred to as wild-type, ΔAHA, and ΔAHA+ pAHA, respectively, in the growth and viability assays described below (**Figures 3A-B, 3E-F**).

### Growth curves

*Hfx. volcanii* strains were grown in liquid Hv-Cab medium at 42 °C with constant orbital shaking until mid-exponential phase (OD_600_ ≈ 0.4) or maintained for an additional 48 h to reach stationary phase (OD_600_ ≈ 4.0). Exponential- and stationary-phase cultures were diluted in fresh Hv-Cab to OD_600_ = 0.05, and 1.5 mL of each culture was transferred into a 12-well flat-bottom plate. Growth curves (≥3 biological replicates per strain) were recorded using an EPOCH2 microplate spectrophotometer (Agilent) at 42 °C with continuous shaking. OD_600_ readings were acquired every 30 min for 48 h. Replicates were averaged and standard deviations calculated.

### Viability assays

Cells were grown to either exponential or stationary phase as described above. Cultures were diluted to OD_600_ = 0.1 in Hv-Cab, serially diluted 10-fold, and 10 µL of each dilution (OD_600_ = 10^−2^ to 10^−6^) was spotted onto Hv-Cab agar plates. Plates were incubated at 45 °C for 2–4 days until colonies formed. Colony-forming units per mL (CFU·mL^−1^) were calculated from countable dilutions. Statistical comparisons were performed using a two-tailed Welch’s t-test on log_10_(CFU·mL^−1^) values.

### Competition assays

Competition experiments were performed by mixing non-fluorescent H26 background strains [wild-type (aBL8) or ΔAHA (aBL638)] with a constitutively fluorescent wild-type-GFP strain (aJM191). Cultures were grown to OD_600_ ≈ 0.4, diluted to OD_600_ = 0.025 in Hv-Cab, and mixed at a 1:1 cell ratio. Mixed cultures were propagated through four consecutive growth–dilution cycles at 42 °C with constant shaking. Each cycle consisted of growth to stationary phase (OD_600_ ≈ 4.0; ∼55–60 h) followed by dilution back to OD_600_ = 0.05 to initiate the next cycle.

Samples for microscopy-based quantification were collected at defined timepoints. In cycle 1, cells were sampled immediately after mixing (initial), during late logarithmic phase (1st exponential; OD_600_ ≈ 0.8), and at stationary phase (OD_600_ ≈ 4.0). For cycles 2–4, samples were collected only at stationary phase.

To quantify relative fitness across serial growth cycles, abundance ratios were log-transformed and analyzed as a function of stationary-phase passages. For each competition, the natural logarithm of the ratio of non-fluorescent to fluorescent cells [ln(non-fluorescent / wt-GFP)] was plotted against the number of completed stationary-phase cycles. Linear regression was performed using data from stationary phases of cycles 1–4, with cycle 1 set as t = 0. The slope of the regression was reported as the selection coefficient per stationary-phase cycle. Lag and logarithmic-phase measurements were excluded from fitness calculations, as no selection is expected prior to stationary-phase passage.

Phase-contrast and GFP fluorescence images were acquired on an Echo Revolve 4 upright/inverted microscope using a 100× Apochromat oil-immersion phase objective (NA 1.3) and a 3.2-MP CMOS camera (effective pixel size: 34.5 nm). GFP was excited using a FITC LED filter cube. Non-fluorescent (competition strain) and fluorescent (wild-type-GFP) cells were manually counted to obtain abundance ratios. For each competition and timepoint, at least 500 cells were analyzed.

### Sample preparation for mass spectrometry

Four biological replicates of each *Hfx. volcanii* strain; either wild-type (aBL8), or ΔAHA (aBL638), were grown in liquid Hv-Cab medium at 42 °C with constant orbital shaking until mid-exponential phase (OD_600_ ≈ 0.5) or further incubated for extra 48 h to reach stationary phase (OD_600_ ≈ 4.0). Cultures were harvested by centrifugation, and cell pellets were flash-frozen and stored at −80 °C until processing.

Following Telusma *et al*., proteins were precipitated from cell pellets with a final concentration of 13% (v/v) trichloroacetic acid (TCA) and centrifuged at 14,000 rpm for 30 minutes at 4°C. The sample precipitates were then washed with cold 10% TCA, washed again with cold acetone, dried at room temperature, and redissolved in 40 μL of 100 mM NH_4_HCO_3_ and 5% acetonitrile (ACN). Protein disulfide bonds were reduced using 5 mM DTT at 65 °C for 10 min, followed by protein alkylation using 15 mM iodoacetamide at 30 °C for 30 min in the dark. Overnight protein digestion was carried out using 0.4 μg of trypsin/lysC (Promega) at 37 °C, followed by an additional 0.2 μg of trypsin/lysC for 1 h before peptides were desalted using Thermo Fisher SOLA SPE plates following manufacturers’ protocols. Desalted peptides were subsequently dried in a speedvac and stored at -80 °C.

### Liquid chromatography–tandem mass spectrometry

LC–MS/MS analysis was performed largely as described previously (Telusma *et al*., 2025). Briefly, tryptic peptides were resuspended in 4% acetonitrile (ACN) and 0.1% formic acid (FA) and separated using an Ultimate 3000 ultra-high-performance liquid chromatography (UHPLC) system (Thermo Fisher Scientific). Approximately 1–2 μg of peptides were loaded onto a C18 trap column (75 μm inner diameter × 2 cm, 3 μm particle size, 100 Å pore size; Thermo Fisher Scientific, #164946) at a flow rate of 5 μL/min and subsequently resolved on a C18 analytical column (75 μm inner diameter × 50 cm, 2 μm particle size, 100 Å pore size; Thermo Fisher Scientific, #ES903). Peptides were washed on the trap column in loading buffer and eluted onto the analytical column using a 135 min linear gradient from 4% to 30% buffer B (ACN, 0.1% FA) at a flow rate of 300 nL/min. Eluting peptides were ionized by nanospray electrospray ionization and analyzed on an Orbitrap Exploris 480 mass spectrometer (Thermo Fisher Scientific).

For data-dependent acquisition (DDA), full MS (MS1) scans were acquired over an m/z range of 390–1390 at a resolution of 120,000, with an automatic gain control (AGC) target of 3 × 10^6^ and a maximum injection time (maxIT) of 50 ms. The top 20 most abundant precursor ions were selected for fragmentation by higher-energy collisional dissociation (HCD) using 25% normalized collision energy (NCE). MS2 spectra were acquired at a resolution of 15,000 with an isolation window of 2.0 m/z, AGC target of 3 × 10^5^, and maxIT of 100 ms.

For data-independent acquisition (DIA), MS1 scan parameters matched those used for DDA. MS2 spectra were acquired using 10 m/z–wide isolation windows with 2 m/z overlaps, spanning the 390–1390 m/z range. Fragment ions were generated using 28% NCE and analyzed at a resolution of 30,000 over a fixed scan range of 200–2000 m/z. AGC and maxIT were set to 2000% and 70 ms, respectively.

### Data-dependent and data-independent proteomic data analysis

DDA raw files were converted to mzML format using MSConvert and searched using the Trans-Proteomic Pipeline (TPP) (Deutsch *et al*., 2010) using the Comet search engine (Eng *et al*., 2013) against the *Haloferax volcanii* Uniprot database (taxon ID 2246). Search parameters include a precursor mass tolerance of 15 ppm, a fragment mass tolerance of 0.02 Da, oxidized methionine as a variable modification, and carbamidomethylation of cysteine as a static modification. Peptide-spectrum matches were validated and scored using PeptideProphet and iProphet (Shteynberg *et al*., 2011). Validated pepXML files were used to generate a spectral library in Skyline (MacLean *et al*., 2010). DIA data were searched against the generated spectral library for extraction of MS1 and MS2 chromatographic peak areas. Peptides corresponding to AHA were manually inspected and curated in Skyline based on peak shape and area, retention time consistency, and signal intensity. MS1-based quantification reports were exported from Skyline and further processed using Python. AHA peptides were normalized to the median intensity of all detected peptides within each sample. Reported AHA protein abundance corresponds to the median abundance across all assigned AHA peptides (n=4).

Following protocols described in Cui *et al*., proteome-wide quantitative analysis was performed using Spectronaut version 16 (Bruderer *et al*., 2016). Briefly, spectral libraries were generated using the DirectDIA workflow with the *Hfx. volcanii* proteome (taxon ID 2246) as the reference database. Default Pulsar and DIA analysis parameters were applied, and quantification was performed at the MS2 level. Differential protein abundance results were exported without filtering, and volcano plots were generated in Python using protein log_2_ fold changes and multiple testing corrected p-values.

### CryoEM sample preparation, grid vitrification and data collection

*Hfx. volcanii* wild-type strain DS2 (aBL3) was grown in Hv-Cab medium to an OD_600_ ≈ 0.75 (late exponential, or the transition from exponential to stationary growth), harvested by centrifugation, and the cell pellets were flash-frozen in liquid nitrogen. For lysate preparation, frozen pellets were resuspended in lysis buffer (20 mM Tris-Cl pH 7.5, 1 M KCl, 10 mM MgCl_2_, 7 mM β-mercaptoethanol, 0.5 mM DTT), lysed by sonication, and clarified by centrifugation. The resulting supernatant was used directly for cryoEM grid preparation.

Grid preparation and vitrification followed a procedure previously described (May *et al*., 2025). Briefly, 300-mesh Quantifoil R2/1 holey-carbon copper grids were coated with a monolayer of graphene and treated with UV/ozone for 10 minutes (Grassetti *et al*., 2023) prior to sample application on the graphene side of the grid. Blotting and plunge-freezing were performed using an FEI Vitrobot Mark IV. Grids were initially screened on Glacios (200 kV) and Tundra (100 kV) microscopes equipped with Falcon III and Falcon-C detectors, respectively (Thermo Fisher Scientific). The final dataset was collected on a Titan Krios G3i operating at 300 kV, using aberration-free image shift (AFIS) in EPU and a Gatan K3 detector mounted on a BioQuantum energy filter. Movies were recorded in super-resolution mode with a total exposure of 50.5 e^−^/Å^2^ fractionated over 50 frames. Data were acquired at a nominal magnification of 81,000X, corresponding to a physical pixel size of 1.06 Å after 2-fold binning. The defocus range was set from –0.2 to –2.4 µm in 0.2-µm increments. In total, 9,140 movies were collected.

### CryoEM data processing and refinement

Movies were processed in CryoSPARC v4.5.3 (Punjani *et al*., 2017) as depicted in **Figure S1**. Briefly, patch motion correction and patch CTF estimation were performed to correct beam-induced motion and to obtain per-micrograph CTF parameters. Micrographs were curated based on CTF fit resolution, defocus range, defocus tilt, relative ice thickness, and total motion; 8,844 of 9,140 movies (96.8%) were retained. Ribosomal particles were identified using reference-free blob picking (particle diameter 170–250 Å; 440-pixel box size). After inspection, ∼1.1 million particles were extracted at 3.64 Å/pixel (128-pixel box). Heterogeneous refinement was then performed using *E. coli* 70S, 50S, and 30S ribosomes plus three junk volumes as templates to accelerate separation of ribosomal and non-ribosomal populations. The 70S class (∼242,000 particles) was retained. To further purify the dataset, iterative *ab initio* reconstruction with 3 classes was performed until all classes produced 70S-like volumes, yielding ∼238,000 particles. These were re-extracted at full resolution (1.06 Å/pixel; 440-pixel box). Global and local CTF refinement and local motion correction were applied to improve per-particle optical and motion parameters, followed by ab initio reconstruction and homogeneous refinement, yielding a 2.47-Å (Gold-Standard Fourier Shell Correlation - GSFSC) consensus map. To resolve structural heterogeneity, 3D classification without a template (10 classes; 8 Å filter) was performed. Six distinct 70S states were identified: four containing tRNAs and two containing AHA. For each of the six particle subsets, a single-class *ab initio* reconstruction / homogeneous refinement was performed, followed by global and local CTF refinement, reference-based motion correction, and a final homogeneous refinement. Maps reached 2.4–2.8 Å resolution (GSFSC). Maps were not masked or processed as isolated 50S/30S bodies because AHA bridges the intersubunit space, and masking could introduce artifacts at this interface. CryoEM map statistics are provided in **Table S1**.

### Model building

CryoEM maps obtained after data processing were used for atomic model construction. Initial models were generated with ModelAngelo v1.0.12 (Jamali *et al*., 2024). A FASTA file containing all expected *Hfx. volcanii* ribosomal proteins and rRNA was provided as input. Additional unassigned densities lacking prior sequence information were traced without sequence constraints to generate ab initio atomic models with per-residue confidence metrics. Sequences extracted from these ab initio models were queried against sequence databases using BLASTP to identify non-canonical factors bound to *Hfx. volcanii* ribosomes in cell lysates. Predicted structures for each protein were retrieved from the AlphaFold2 database (Fleming *et al*., 2025; Varadi *et al*., 2024) and rigid-body docked into the cryoEM maps using ChimeraX (Meng *et al*., 2023). Protein models were further adjusted and rebuilt manually in Coot (Emsley *et al*., 2010). For rRNA, structures of the closest available archaeal homologues [23S and 5S rRNAs from *Haloarcula marismortui* (PDB: 4V9F) and 16S rRNA from *Thermococcus celer* (PDB: 6TMF)] were docked into the maps and mutated to the *Hfx. volcanii* sequence using Web 3DNA 2.0 (Li *et al*., 2019), followed by iterative fitting in Coot. Final composite models were refined using Phenix real-space refinement (Afonine *et al*., 2018) and validated with MolProbity (Williams *et al*., 2018) to assess geometry, stereochemistry, and model–map agreement. CryoEM refinement statistics and model–map correlation data are provided in Table S1. Structural figures were prepared using ChimeraX (Meng *et al*., 2023). The structures of the *Hfx. volcanii* 70S ribosome representing translational states are not the focus of this work and will be described elsewhere.

### Evolutionary and bioinformatics analyses

The evolutionary distribution and sequence diversity of *Hfx. volcanii* AHA (HVO_2384) were investigated by combining profile HMM-based and structure-based similarity searches. Profile HMM-profile HMM comparisons were performed using HHpred via the MPI Bioinformatics Toolkit (Zimmermann *et al*., 2018), querying the PDB70 database under default parameters. In parallel, structure-based searches were conducted with FoldSeek (van Kempen *et al*., 2024) using default settings.

To identify AHA homologs across archaea, we first performed a single PSI-BLAST (Camacho *et al*., 2009) iteration against the archaeal subset of the non-redundant database (nr_arc) using default parameters. This search predominantly retrieved close homologs within Halobacteriota, suggesting limited sensitivity toward more divergent archaeal representatives. We therefore complemented these analyses with additional HHpred searches against archaeal proteomes and FoldSeek comparisons, which recovered more distant candidates across multiple archaeal lineages. Based on these results, representative HPF-domain sequences from major archaeal phyla were selected as independent PSI-BLAST seeds to systematically expand the dataset. The HPF domain was used because it represents a defining and conserved feature of AHA, whereas CBS domains are widespread among diverse protein families and are not specific to AHA. The following sequences were used as queries: *Hfx. volcanii* (UniProt D4GWN2), *Thermofilum pendens* (A1S180), *Saccharolobus solfataricus* (A0A0E3MD01), *Nitrososphaera viennensis* (A0A977ICN8), *Methanomassiliicoccales archaeon RumEn M2* (A0A0Q4BAF9), *Candidatus Nanohalobium* (A0A5Q0UFT2), *Fermentimicrarchaeum limneticum* (A0A7D6BN20), *Pyrococcus furiosus* (A0A5C0XMJ8), and *Methanopyrus kandleri* (Q8TZ50). Searches were performed against the GTDB archaeal protein dataset (release RS226; 6,968 genomes) (Parks *et al*., 2025), which provides a phylogenetically standardized and taxonomically comprehensive framework for assessing distribution across archaeal diversity. PSI-BLAST searches were run for two iterations with an inclusion threshold and E-value cutoff of 1e^−5^. All hits from the different query seeds were pooled and filtered to remove sequences shorter than 70 amino acids and entries annotated as partial (partial=01, 10, or 11), yielding 4,107 sequences. The dataset was aligned using MAFFT (Katoh & Standley, 2013) in automatic mode and inspected to distinguish canonical AHA proteins, characterized by an N-terminal 4×CBS module followed by a C-terminal HPF domain, from proteins containing only the HPF domain. Sequences failing to align coherently across the HPF domain or CBS region were removed. The final curated archaeal HPF dataset comprised 3,160 AHA proteins and 926 HPF-only proteins. Their phylogenetic distribution was mapped onto the GTDB RS226 archaeal reference tree using the iTOL webserver (Letunic & Bork, 2024).

To explore the sequence space of AMPKγ homologs and evaluate their potential evolutionary relationship with AHA, we used the PANTHER (Thomas *et al*., 2022) AMPKγ profile HMM (PTHR13780) as query. HMMER searches were performed against UniProt and supplemented with GTDB archaea to ensure comprehensive archaeal coverage. Hits were retained using an E-value cutoff of 1.0 × 10^−5^ and a minimum HMM coverage threshold of 60%. Retrieved full-length sequences were pooled and clustered with MMseqs2 at 70% sequence identity and 70% coverage, resulting in a CBS dataset containing 5,820 representative sequences. Sequence relationships were analyzed using CLANS with a stringent cutoff of 1.0 × 10^−13^.

With the aim of assessing the phylogenetic distribution of SriA-SriD and Dri across archaea, homologs were retrieved from GTDB using PSI-BLAST with an E-value cutoff of 1.0 × 10^−5^. Queries included SriA–SriD from *Methanosarcina acetivorans C2A* (Q8TH74, Q8TH73, Q8TH72, Q8TH71) and Dri from *Pyrobaculum calidifontis* (A3MX58). Only hits longer than 200 amino acids were retained, and sequences annotated as partial (partial=01, 10, or 11) were excluded to avoid truncated or incomplete entries. Homologs of SriA–SriD were pooled (13,735 sequences) and analyzed using CLANS based on all-against-all BLAST P-values to resolve the four subfamilies in sequence space. For all protein families analyzed, structural models were retrieved from the Protein Data Bank (wwPDB consortium, 2018) and AlphaFold database (Varadi *et al*., 2024) where available; sequences lacking solved or precomputed models were predicted using the AlphaFold3 server (Abramson *et al*., 2024).

For evolutionary analysis of the HPF domain, the archaeal HPF dataset (above) was extended to include bacterial and eukaryotic homologs. Bacterial representatives were retrieved using *E. coli* HPF (P0AFX0), *E. coli* RaiA (P0AD49), and *Bacillus subtilis subsp. subtilis* HPF (A0A1B2AZA1) as PSI-BLAST seeds against GTDB Bacteria and UniProt (UniProt Consortium, 2024). All bacterial and eukaryotic hits were combined with the archaeal HPF sequences to generate a cross-domain dataset. Redundancy was reduced using MMseqs2 (Steinegger & Soding, 2017) clustering at 80% sequence identity and 70% coverage, resulting in 10,581 representative sequences. Global sequence relationships were examined using CLANS (Frickey & Lupas, 2004; Zimmermann *et al*., 2018), based on all-against-all BLAST P-values (cutoff 1.0 × 10^−6^). For phylogenetic reconstruction, 500 representative HPF sequences each from archaea and bacteria were selected using taxonomy-balanced sampling across phyla and families, preferentially choosing sequences close to the median length within each family. Archaeal sequences spanned 18 of the 21 archaeal phyla currently recognized by the Genome Taxonomy Database (GTDB) (Parks *et al*., 2025), while bacterial sequences were sampled from 170 of the 183 GTDB-defined bacterial phyla, representing approximately 86% and 93% of archaeal and bacterial phylum-level diversity, respectively.

For the generation of the HPF phylogenetic tree, sequences were trimmed to the conserved HPF domain prior to alignment. Multiple sequence alignment was performed using MAFFT v7.526 with the L-INS-i algorithm (local pair with iterative refinement) (Katoh & Standley, 2013), which is appropriate for divergent single-domain proteins. Poorly aligned and sparsely occupied positions were detected and removed using trimAl v1.5 in gappyout mode (Capella-Gutiérrez *et al*., 2009), yielding a final alignment of 127 amino acid positions.

Maximum-likelihood phylogenetic inference was carried out using IQ-TREE3 v3.0.1 (Wong *et al*., 2025), with automatic model selection implemented in ModelFinder (Kalyaanamoorthy *et al*., 2017). The best-fit substitution model was selected according to the Bayesian Information Criterion (BIC). Branch support was assessed using 1,000 ultrafast bootstrap replicates (UFBoot2, with branch-length optimization) (Hoang *et al*., 2017) and 1,000 SH-like approximate likelihood ratio test (SH-aLRT) replicates (Guindon *et al*., 2010). Phylogenetic trees were visualized and rendered using FigTree v1.4.4 (Rambaut, 2018).

To assess the robustness of the inferred topology, phylogenetic analyses were repeated using alternative alignment and trimming strategies. Additional alignments were generated using MAFFT’s automatic alignment strategy, and trimming was performed using a more stringent gap-threshold filter (trimAl -gt 0.5). Across all analyses, model selection was consistent, with Q.PFAM+F+G4 identified as the best-fit model according to both the Bayesian and Akaike information criteria. The Q.PFAM+F+G4 model accounts for amino acid exchange rates derived from Pfam domain alignments (Q.PFAM), empirical amino acid frequencies (+F), and across-site rate variation modeled using a discretized Gamma distribution with four rate categories (+G4).

For phylogenetic analyses of CBS domains, up to 50 representative sequences per major homologous cluster were selected from the CBS dataset (above) using taxonomy-balanced sampling, for a total of 637 CBS sequences. Non-CBS regions were removed prior to multiple sequence alignment, which was performed with MAFFT v7.526. Several alignment strategies (L-INS-i, E-INS-i, and auto) were evaluated; L-INS-i was selected because it produced a compact alignment with higher column occupancy.

To minimize noise from poorly aligned or sparsely populated positions, multiple trimming strategies were tested in trimAl v1.5, including -gappyout, -automated1, and fixed gap thresholds (-gt 0.5, 0.6, 0.7, 0.8, 0.9, and 0.95). Maximum-likelihood trees were inferred for each trimmed alignment using IQ-TREE3 v3.0.1. Model selection was performed with ModelFinder (-MFP, extended search), which consistently identified LG+R9 as the best-fitting model according to both Bayesian (BIC) and Akaike (AIC) information criteria. The LG+R9 model combines the LG amino acid empirical substitution matrix with across-site rate heterogeneity modeled using the FreeRate approach (+R9), which estimates nine rate categories and their proportions directly from the data. Branch support was assessed using 1,000 ultrafast bootstrap replicates and 1,000 SH-aLRT replicates. As tree topologies were consistent across trimming strategies, the -gt 0.6 alignment (254 amino acid positions) was selected as a representative phylogeny for downstream analyses.

## ACKNOWLEDGEMENTS

We thank Sarah Sterling, Jenn Podgorski, Nicholas Swanson, and Dan Cham-Chin Lim from the MIT.nano CryoEM facility, and Berith Isaac from the Brandeis University EM facility. CryoEM specimens were prepared and data were collected at the CryoEM facility in MIT.nano, including use of the Titan Krios and Talos Arctica, the latter gifted by the Arnold and Mabel Beckman Foundation. CryoEM screening data were also collected at the Brandeis EM facility using a Tundra Cryo TEM, purchased under the NSF-DBI 2319804 award. We thank John Mallon for generating the wild-type-GFP (aJM191) strain and Zachary Curtis for the cloning of the pAL750-AHA (eZC128) plasmid. This work was supported by grants NIH R01-GM144542 (JHD), T32-GM136540 (MBM), and R35-GM156992 (AB), NSF CAREER-2046778 (JHD), MCB-2222076 (AB), and GRFP (MBM), the Max Planck Society (VA), and HFSP-RGY0074 (AB and VA). AB is a Pew Scholar in the Biomedical Sciences, supported by The Pew Charitable Trusts. The authors used AI-assisted language tools (ChatGPT, OpenAI) for grammatical and stylistic copy-editing; all scientific content was written, verified, and is the sole responsibility of the authors.

## AUTHOR CONTRIBUTIONS

D.P.S., A.B., and J.H.D. conceived the project. D.P.S performed all cell-based experiments, prepared samples, acquired and processed cryoEM data, built atomic models, and conducted the phylogenetic analyses. M.B.M. carried out the mass spectrometry experiments and assisted with cryoEM grid and sample preparation, and data collection. J.C contributed to ModelAngelo-based model building and supported cryoEM data processing. V.A. performed the HPF and CBS evolutionary clustering analyses and examined the phylogenetic distribution of these domains across archaea. A.B. and J.H.D. supervised the study and secured funding. D.P.S. wrote the first draft of the manuscript, with contributions, feedback, and editing from all authors.

## DATA AND MATERIALS AVAILABILITY

Upon publication, raw micrographs will be deposited at EMPIAR, and atomic coordinates and the cryoEM map of the *Hfx. volcanii* ribosome bound to AHA and E-site tRNA will be deposted at the Protein Data Bank (PDB) and Electron Microscopy Data Bank (EMDB), respectively. Strains are available upon request.

