## Supplementary Information for "A conserved archaeal ribosome-associated factor linking bacterial hibernation and eukaryotic energy sensing"

### SUPPLEMENTARY TABLES

| Data acquisition | | | | | | |
|---|---|---|---|---|---|---|
| Microscope | Titan Krios G3i | | | | | |
| Camera | Gatan K3 (counting mode) | | | | | |
| Magnification (X) | 81,000 | | | | | |
| Voltage (kV) | 300 | | | | | |
| Total electron dose ($e^-/Å^2$) | 50.51 | | | | | |
| Defocus range (µm) | -0.2 to -2.4 (step size: 0.2) | | | | | |
| Pixel size (Å) | 1.06 | | | | | |
| Micrographs collected (#) | 9,140 | | | | | |
| **Reconstruction** | | | | | | |
| **Reconstruction** | **AHA + E/E-tRNA** | **AHA alone** | **P/P-tRNA** | **A/A-, P/P-tRNA** | **P/P- , E/E-tRNA** | **A/P-, P/E-tRNA** |
| Processing suite | cryoSPARC | | | | | |
| Symmetry | C1 | | | | | |
| Final particle count | 82,088 | 70,830 | 27,005 | 17,522 | 27,489 | 12,771 |
| Resolution (Å) (GS-FSC (0.143) | | | | | | |
| *masked (tight, corrected)* | 2.4 | 2.4 | 2.6 | 2.7 | 2.6 | 2.8 |
| *unmasked* | 3.1 | 3.1 | 3.5 | 3.9 | 3.6 | 4.3 |

**Table S1.** CryoEM data collection and processing information.

| Strain | Genotype | Reference |
|---|---|---|
| **DS2 (aBL3)** | - | Hartman *et al.*, 2010 |
| **H26 (aBL8)** | ΔpyrE2 | Allers *et al.*, 2004 |
| **ΔAHA (aBL638)** | ΔpyrE2 ΔHVO_2384 | This work |
| **H26 pAL750 (aBL639)** | ΔpyrE2 pAL750 | This work |
| **ΔAHA pAL750 (aBL640)** | ΔpyrE2 ΔHVO_2384 pAL750 | This work |
| **ΔAHA pAL750-AHA (aBL641)** | ΔpyrE2 ΔHVO_2384 pAL750::HVO_2384 | This work |
| **H26-GFP (aJM191)** | ΔpyrE2 pyrE2::p27-msfGFP-mevR | This work |

| Plasmid | Genotype (description) | Reference |
|---|---|---|
| **pAL750 (eTR37)** | pAL750 (empty vector; Pxyl pro-moter) | (Rados et al., 2023) |
| **pAL750-AHA (eZC128)** | pAL750::HVO_2384 (AHA expres-sion under Pxyl) | This work |
| **pTSD1 (eTR68)** | pTSD1 (knock out generation in Hfx. volcanii) | (Rados et al, 2025) |
| **pTSD1-AHA (eDS1)** | pTSD1::HVO_2384u-HVO_2384d (ΔAHA generation) | This work |
| **pBlue-GFP (eJM53)** | pBlue-pyrE2::p27-msfGFP-mevR (msfGFP expression) | This work |

| Primers | Sequence (5′→3′) | Use |
|---|---|---|
| **pTSD1_HVO_2384-up-fw** | CCGAAAAGTGCCACCTTCGTCGAAGACGACGTTC | eDS1 generation |
| **HVO_2384-up-rv** | CATCCTTACATACAACGCCG | eDS1 generation |
| **HVO_2384-up-HVO_2384-down-fw** | CGGCGTTGTATGTAAGGATGCAAGCTCAACGAGCTGTAAG | eDS1 generation |
| **pTSD1_HVO_2384-down-rv** | CGAGGGGTTTTATCCACGGTGGCGACGACTATCAC | eDS1 generation |
| **Pxyl_HVO_2384-fw** | GCTGGTAATGAGGATACTGCAATGGATATTGCTGATATCGCC | eZC128 generation |
| **Pxyl_HVO_2384-rv** | GGCCGCTCTAGAACTAGTTTACAGCTCGTTGAGCTTG | eZC128 generation |
| **pBlue-pyrE2-up-fw (oBL136)** | CCGAAAAGTGCCACCTCGGCGGTAGAAGTACG | eJM53 generation |
| **pyrE2-up-rv (oBL87)** | CTTGTTTCGAGAGGGTTCAG | eJM53 generation |
| **p27_pyrE2-up-fw (oJM212)** | CTGAACCCTCTCGAAACAAGACCCGCCGACTCGGCGT | eJM53 generation |
| **msfGFP_mevR-rv (oJM213)** | TCTTCCGCTTCCTCGTCATTTGTAAAGTTCATCCATTCCAT | eJM53 generation |
| **pyrE2-mevR-fw (oBL88)** | CTGAACCCTCTCGAAACAAGCGAGGAAGCGGAAGAG | eJM53 generation |
| **mevR_rv_pyrE2 (oJM160)** | AGTGAATTAGTCACAAGCGTGTACATGGGAGGGGATGG | eJM53 generation |
| **pyrE-down-fw (oHV42)** | TACACGCTTGTGACTAATTCACT | eJM53 generation |
| **pyrE2-down-rv (oBL137)** | GCTGGCCTTTTGCTCAGTGGCGGATTCGATGTAG | eJM53 generation |
| **Pscript-fw (oBL105)** | TGAGCAAAAGGCCAGC | eJM53 generation |
| **pTA131-Pamp-rv (oBL97)** | AGGTGGCACTTTTCGG | eJM53 generation |

**Table S2.** Strains, plasmids and primers used in this work.

### SUPPLEMENTARY FIGURES

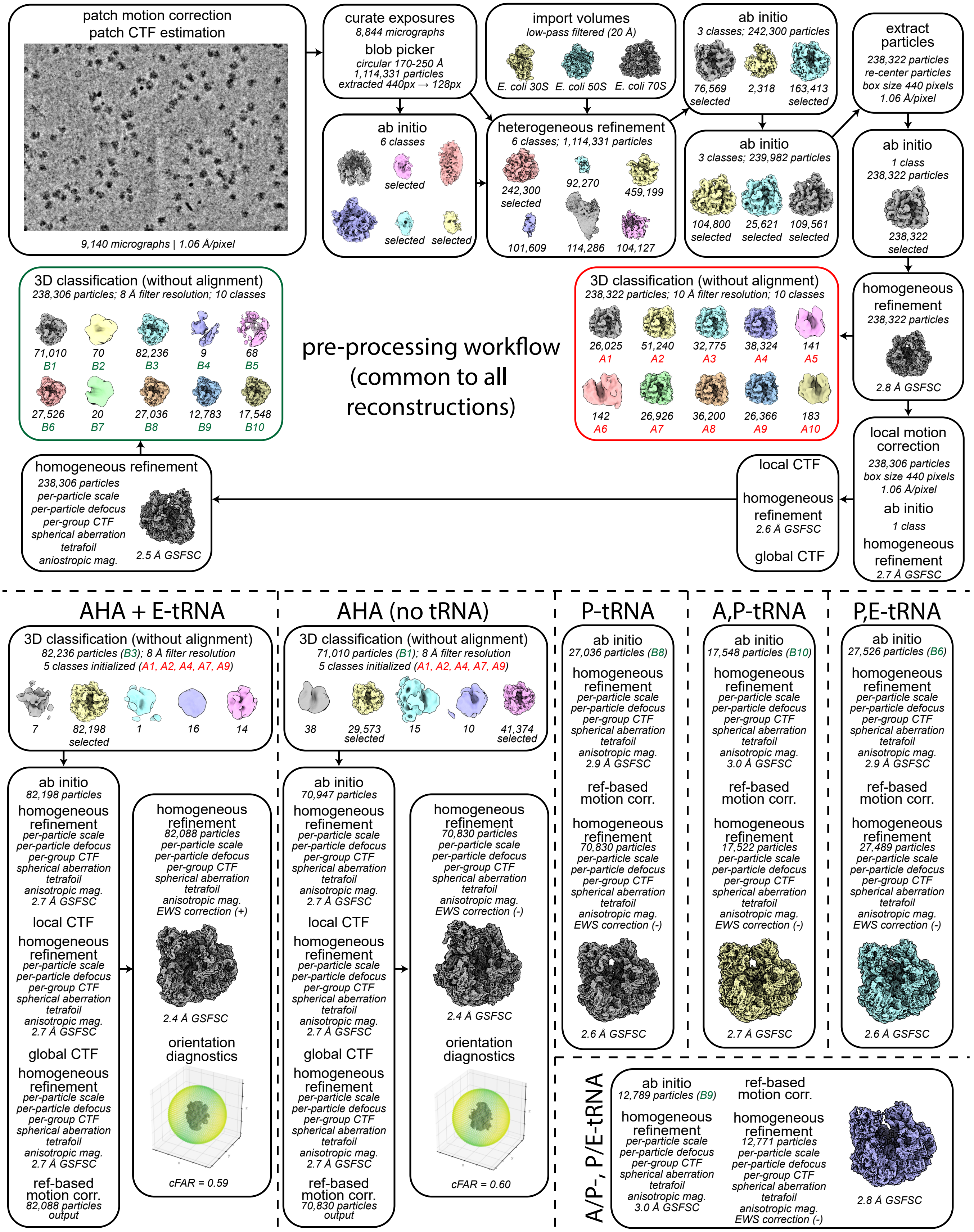


**Figure S1. CryoEM image processing workflow.** Schematic of the cryoSPARC-based (Punjani et al., 2017) cryoEM data processing pipeline used to obtain high-resolution consensus 70S ribosome reconstruction (top) and to classify relevant substates (bottom). Job names, particle counts, key metadata, and any non-default parameters are noted.

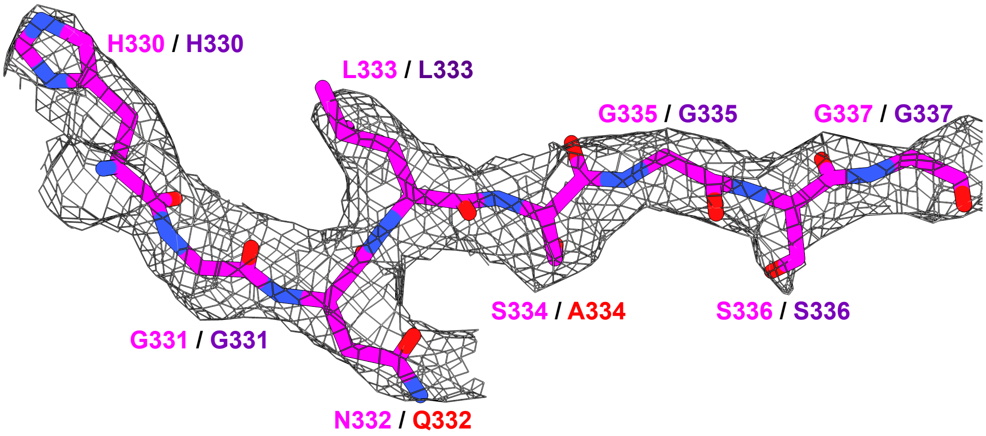


ModelAngelo 330 HGNLSGSG 337
HG L+GSG
Hvo_2384/AHA 330 HGQLAGSG 337

**Figure S2. Automated atomic modeling-based identification of HVO_2384 (AHA) bound to a translationally inactive ribosome.** CryoEM density (black mesh) from the AHA + E/E-site tRNA map (Figure 1F, Table S1) is shown with the unrefined, ModelAngelo-built atomic model docked into the map (top). Below, the amino acid sequence inferred by ModelAngelo (pink) is aligned with the annotated HVO_2384 sequence in this region (purple). Residue mismatches (highlighted in red) reflect positions where amino acids with chemically similar side-chains cannot be unambiguously distinguished at this resolution when the sequence is not provided to the algorithm.

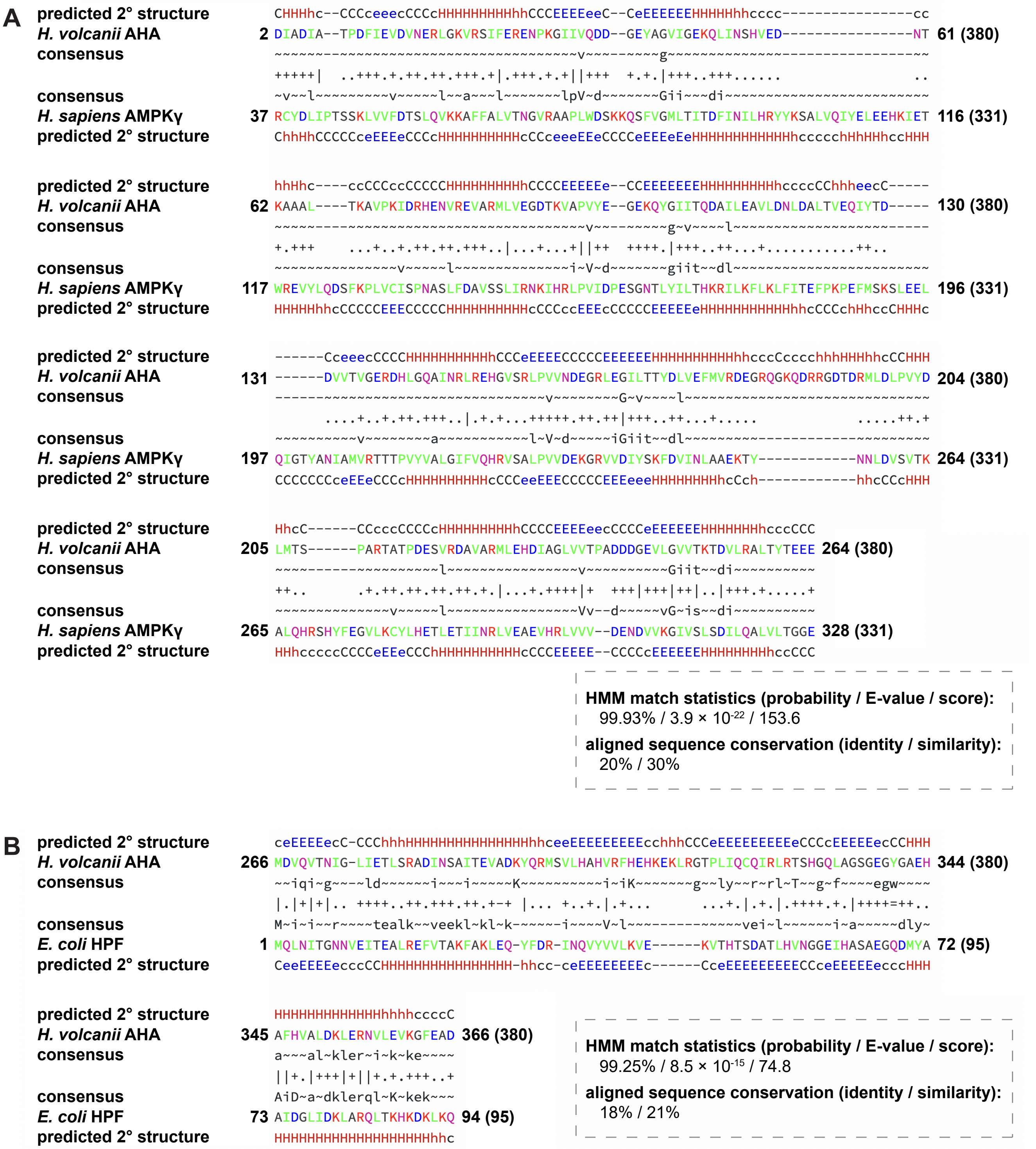


**Figure S3. Archaeal AHA comprises an N-terminal CBS tetrad homologous to eukaryotic AMPKγ and a C-terminal HPF-like domain homologous to bacterial HPF.** Homology between the N- and C-terminal domains of *Hfx. volcanii* AHA (HVO_2384) and **(A)** human AMPKγ (Uniprot: P54619) and **(B)** *E. coli* HPF (Uniprot: P0AFX0) was identified using HHpred (Zimmermann *et al.*, 2018), based on profile–profile comparison of their respective hidden Markov models (HMMs). Shown are the optimal HHpred alignments, with probabilities, E-values and scores indicated. Despite low pairwise sequence identity (20% for CBS-AMPKγ and 18% for HPF), the high HMM probabilities support deep evolutionary relationships. For the CBS comparison, the AHA N-terminal sequence was used to generate an HMM profile that was searched against a *Homo sapiens* profile database. For the HPF comparison, HMM profiles were constructed from provided multiple sequence alignments of archaeal AHA homologues (64 proteins) and bacterial HPF proteins (53 proteins). Residue numbering is indicated at the start and end of each sequence block. Consensus rows represent the most probable residue at each HMM position; uppercase and lowercase letters reflect the strength of conservation in the profile. Alignment symbols denote similarity inferred from the profile–profile comparison: | identical residues; + strong similarity; . weak similarity; blank, no detectable similarity. Amino acids are colored by physicochemical class (apolar, green; positively charged, red; negatively charged, blue; polar uncharged, magenta; small residues, black). Predicted secondary structure is shown above and below the sequences (helices, H/h, red; β-strands, E/e, blue; coils C/c, black; uppercase indicates higher confidence).

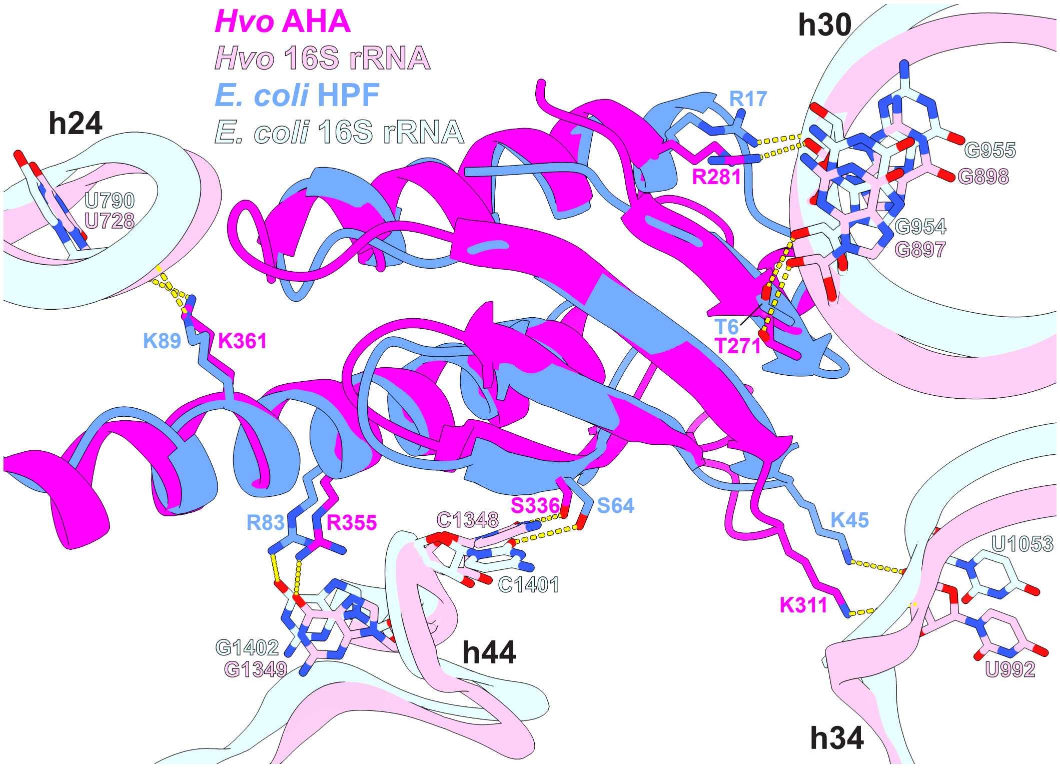


**Figure S4. Conserved ribosomal recognition by archaeal and bacterial HPF domains.** Structural comparison of archaeal (*Hfx. volcanii*, this work) and bacterial (*E. coli*, PDB 6Y69) HPF domains bound to the small ribosomal subunit highlighting a conserved set of interactions with 16S rRNA. Archaeal and bacterial HPFs and 16S rRNAs are colored following the legend. Equivalent residues that contact conserved 16S rRNA residues in each system are indicated, together with the corresponding 16S rRNA helix numbers.

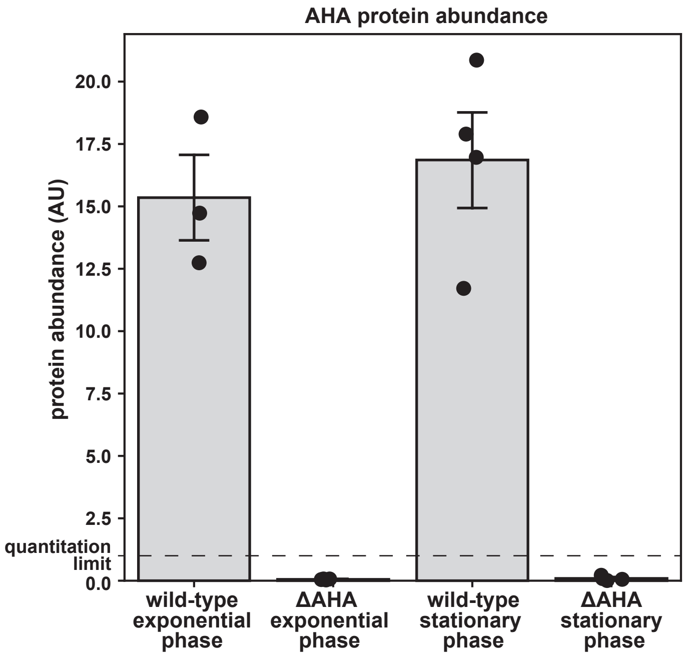


**Figure S5. AHA expression is not regulated by growth phase.** Quantitative mass spectrometry analysis of AHA protein abundance in wild-type and ΔAHA *Hfx. volcanii* cells during exponential and stationary phase growth. AHA peptides (n = 4) were robustly detected and quantified in wild-type cells at similar levels in both growth states, whereas AHA-derived peptides in the ΔAHA strain remained below the quantitation limit of the assay (dashed line). Data points represent individual biological replicates (n≥3); bars denote mean abundance ± standard error of the mean.

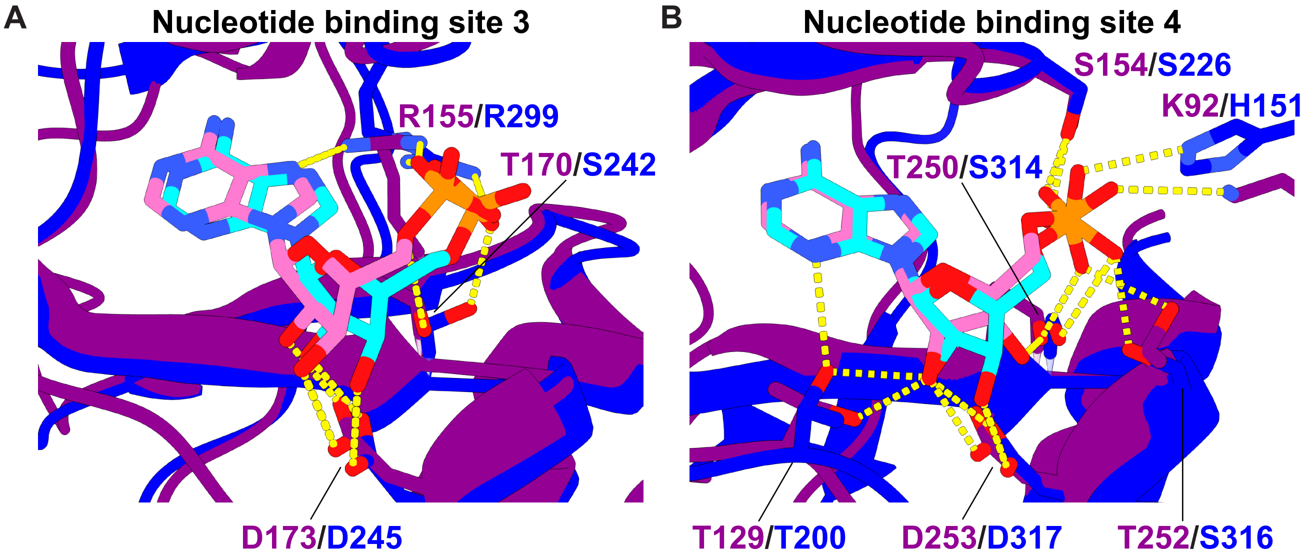


**Figure S6. Comparison of the nucleotide binding sites in AHA and AMPKγ.** Structural superposition of the AMP-binding pockets in AHA 4×CBS domain (protein chain and amino acid labels in purple; AMP carbon atoms in pink) and the eukaryotic AMPKγ subunit (PDB: 6B1U), with its protein and amino acid labels in dark blue, and AMP molecules in cyan. Equivalent or similar protein side chains forming hydrogen bonds or electrostatic interactions with the nucleotides are labeled, and interactions are indicated by yellow dashed lines. AHA's arginine (R) 155 and AMPKγ's R299 are not homologous residues as they are in different β-strands but similarly interact with the same AMP region. Oxygen, nitrogen, and phosphorus atoms are colored red, blue, and orange, respectively.

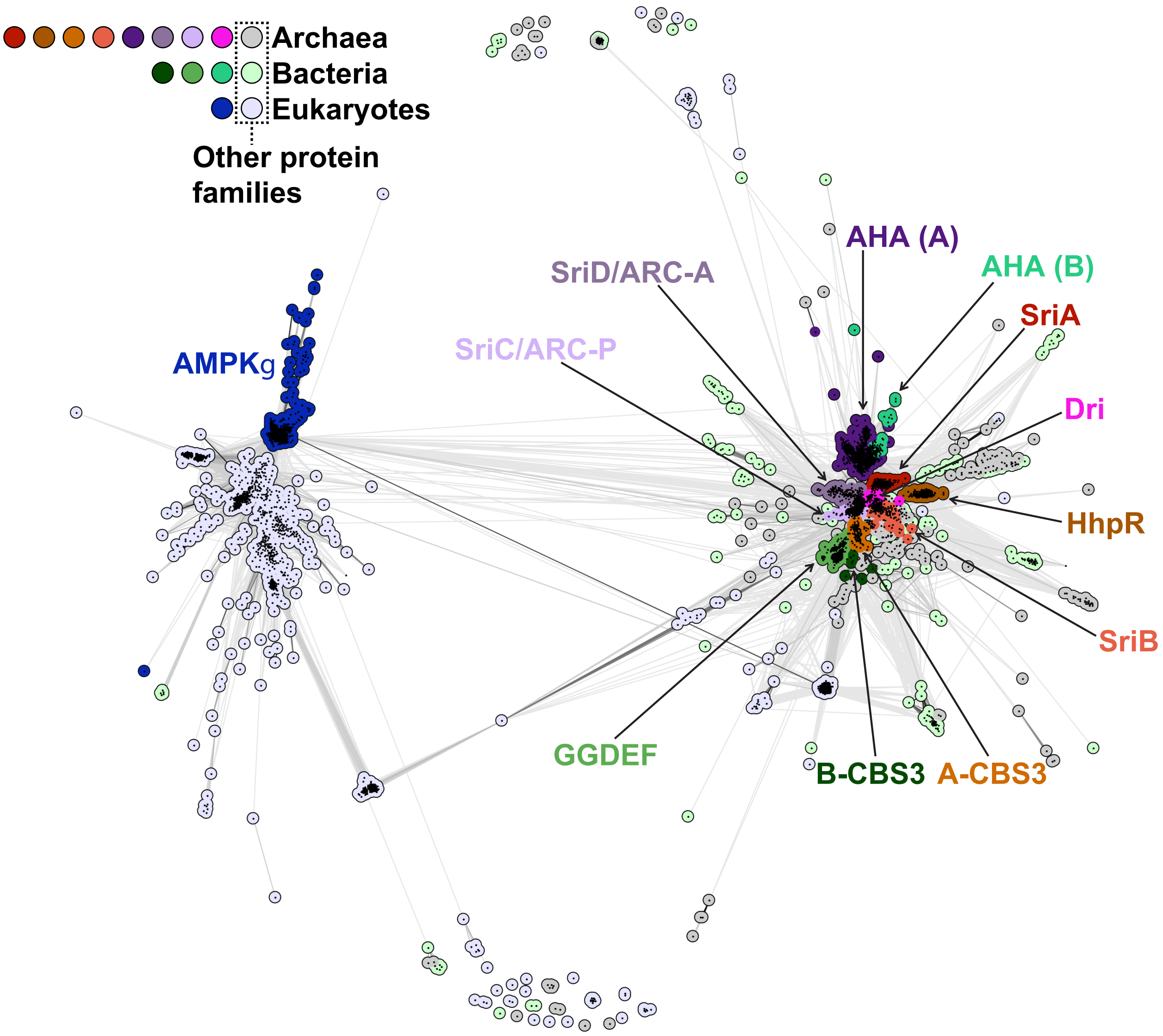


**Figure S7. CLANS cluster-map analysis of the CBS-domain superfamily based on all-against-all pairwise sequence similarities.** Each node represents an individual protein, and connecting lines reflect BLAST p-values, with darker lines indicating higher sequence similarity. Proteins are colored by taxonomic origin: bacteria (shades of green), eukaryotes (dark blue and lavender-blue), and archaea (all other colors). Protein families labeled: archaeal (A) AHA, Dri, SriA, SriB, SriC/ARC-P, SriD/ARC-A, HhpR and A-CBS3; bacterial (B) AHA homolog, GGDEF-containing 4×CBS fusion (GGDEF) and B-CBS3; eukaryotic AMPKγ.

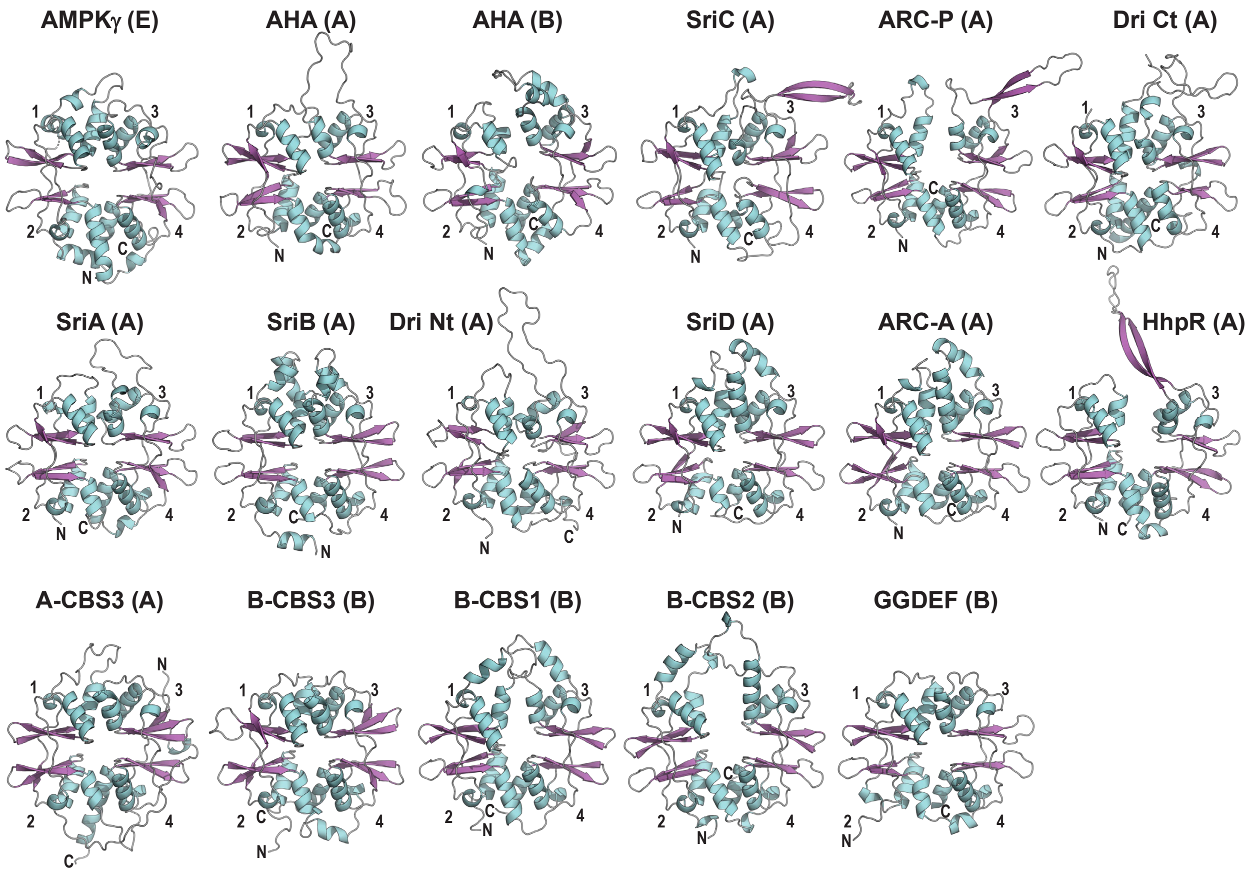


**Figure S8. Structural comparison of members of the CBS-domain protein superfamily related to the N-terminal region of AHA.** Representative CBS-containing proteins from eukaryotes (E), archaea (A), and bacteria (B) are shown [species; structural source (PDB or AlphaFold Database, AFDB) accession]: AMPKγ (Homo sapiens; PDB: 6B1U); AHA (*Hfx. volcanii*; cryoEM structure, this work); bacterial AHA homolog (*Candidatus Gottesmanbacteria bacterium*; AFDB: A0A1F5YVL3); SriA, SriC, SriD (*Methanosarcina acetivorans*; PDB: 9ZNF, 9ZNI, 9ZNI); SriB (*Methanosarcina acetivorans*; AFDB: Q8TH73); ARC-P and ARC-A (*Methanolobus sp. ZRKC1*; NCBI: WYM85633.1 and WYM85634.1; computed using AlphaFold3); Dri N-terminal (Nt) and C-terminal (Ct) domains, shown separately (*Pyrobaculum calidifontis*; PDB: 9E6Q and 9E7F); HhpR (*Pyrococcus yayanosii*; AFDB: F8AFU3); archaeal CBS3 (A-CBS3; miscellaneous Crenarchaeota group-1 archaeon SG8-32-3; AFDB: A0A0M0BVG1); bacterial CBS1 (B-CBS1; *Anaerolinea thermophila*; AFDB: E8N4C0); bacterial CBS2 (B-CBS2; *Deinococcus reticulitermitis*; AFDB: A0A1H6XD96); bacterial CBS3 (B-CBS3; *Candidatus Nitrospira nitrificans*; AFDB: A0A0S4LEU7); and GGDEF-containing 4×CBS fusion proteins (*Romeriopsis navalis*; AFDB: A0A928VUH7). The four CBS motifs within each protein (CBS1–CBS4) are labeled numerically. N- and C-termini are indicated for reference. α-helices are shown in teal and β-strands in pink. The long, extended disordered N-terminal region of SriB is omitted for clarity, whereas the N-terminal extension in A-CBS3 is shown.
